# Localized translation of cell junction mRNAs is required for epithelial cell polarity

**DOI:** 10.1101/2024.12.16.628746

**Authors:** Ashley Chin, Jonathan Bergeman, Laudine Communal, Jonathan Boulais, Anne-Marie Mes-Masson, Eric Lécuyer

## Abstract

Epithelial cells exhibit a highly polarized organization along their apico-basal axis, a feature that is critical to their function and frequently perturbed in cancer. One less explored process modulating epithelial cell polarity is the subcellular localization of mRNA molecules. In the present study, we report that several mRNAs encoding evolutionarily conserved epithelial polarity regulatory proteins, including *Zo-1*, *Afdn* and *Scrib*, are localized to cell junction regions in *Drosophila* epithelial tissues and human epithelial cells. Targeting of these mRNAs is coincident with the robust junctional distribution of their encoded proteins, and we demonstrate that they are locally translated at cell junction regions. To identify RNA binding proteins (RBPs) potentially implicated in junctional mRNA regulation, we performed systematic immuno-labeling with a collection of validated RBP antibodies, identifying a dozen RBPs with consistent junctional distribution patterns, several of which directly bind junctional transcripts. Strikingly, depletion of these RBP candidates, including MAGOH, a core component of the exon-junction complex (EJC), perturbed the junctional distribution and localized translation of *Zo-1* and *Scrib* mRNAs, as well as the junctional accumulation of their protein products. Functional disruption of MAGO, or its interaction partner Y14, in *Drosophila* follicular epithelial cells perturbs the distribution of junctional transcripts and proteins. Finally, tissue microarray analysis of ovarian cancer tumor specimens revealed that expression of MAGOH and ZO-1 is positively correlated and that both proteins are potential biomarkers of good prognosis. Altogether, this work reveals that localized mRNA translation at cell junction regions is important for modulating epithelial cell polarity.

**HIGHLIGHTS:**

- Cell junction mRNA targeting is conserved between tissues and species
- These mRNAs undergo localized translation at areas of cell-cell contact
- A diversity of RBPs localize to cell junction regions and interact with junctional transcripts.
- Disruption of junctional RBPs impacts epithelial cell polarity and localized translation
- MAGOH and ZO-1 expression is correlated in ovarian tumor specimens and are potential biomarkers of good prognosis

## INTRODUCTION

Cells come in diverse shapes and sizes, often presenting an asymmetrical organization that is adapted to their function. This is exemplified by epithelial cells that compose the cellular sheets that shield our tissues from the external environment and are highly polarized along their apico-basal axis [1]. Epithelial cell polarization relies on the presence and function of cell junction structures, including adherens and tight junctions, which serve as mediators of cell-cell adhesion and selective permeability barriers [2]. This polarity is coordinated by evolutionarily conserved protein modules, including transmembrane and cytosolic adaptor proteins that make up the cell junctions, as well as polarity complexes that demarcate specialized membrane territories along the apico-basal axis of the cell [3–5]. Importantly, loss of cell polarity is a hallmark of carcinomas, malignancies of epithelial origin that encompass the majority of human cancers [5, 6]. Indeed, perturbed expression and localization of polarity regulators has been linked to poor prognosis in cancer patients, while genetic studies in *Drosophila* and mice identified these factors as neoplastic tumor suppressors, whereby their inactivation disrupts polarity and contributes to tumorigenesis [7–16]. Thus, defining the mechanisms that regulate the establishment and maintenance of epithelial cell polarity is a key biological question [17–19].

An emerging mechanism implicated in the functional organization of epithelial cells is the asymmetric subcellular distribution of RNA molecules [20, 21]. While the intracellular localization of proteins can be modulated via co-or post-translation mechanisms, following a traditional signal peptide model, the trafficking and localized translation of mRNA offers an exquisite strategy to target proteins on-site and on-demand, while preventing dispersion to non-productive sites in the cell [22]. In the context of epithelial cells, the localization of mRNAs such as *Crumbs, Wingless, Stardust* and *Zonula occludens 1* (*ZO-1*) to the apical domain of *Drosophila* and mouse epithelia was found to be important for accurate protein targeting, morphogenetic signaling and cell polarity modulation [23–26]. Moreover, large-scale transcriptome mapping efforts have revealed the prevalent asymmetric distribution properties of RNA molecules in epithelial cells and tissues [27–29]. For example, systematic fluorescent *in situ* hybridization (FISH)-based surveys in fly embryos identified a few dozen mRNAs localized asymmetrically along the apico-basal axis and lateral membranes of embryonic epithelial cells, many of which encode evolutionarily conserved cell junction components such as Canoe (CNO), Discs Large-1 (DLG1), Scribble (SCRIB), and Polychatoid (PYD)/*Drosophila* ZO-1 (dZO-1) [27, 30]. Transcriptomic profiling of micro-dissected apical and basal compartments of mouse intestinal epithelia recently revealed a high prevalence for polarized RNA distribution [28]. Strikingly, the authors observed that apically-enriched mRNAs are translated more efficiently owing to the co-enrichment of ribosomes in the apical cytoplasm, a process that is dynamically regulated to enhance nutrient absorption during feeding. It was also recently found that PLEKHA7, a protein localized to adherens junctions, can help recruit core components of the RNA-induced silencing complex (RISC) and microRNA molecules to cell junctions regions [31, 32], implicating local miRNA-based regulation at cell junction structures.

In the present study, we demonstrate that several mRNAs encoding cell junction proteins (e.g. *Zo-1, Scrib and Afdn*) are targeted to junctional regions of epithelial cells, both in *Drosophila* tissues and human cell lines, and that these transcripts undergo localized translation. To identify potential regulators of junctional mRNAs, we performed imaging-based screens in human epithelial cell models, using a validated collection of antibodies directed against hundreds of RBPs, which identified a dozen factors with consistent cell junction patterns. We found that RBPs localized to the cell membrane regions robustly associate with junctional mRNA candidates. Functional assays revealed that several of these RBPs are required for the proper targeting of junctional proteins and for the localized translation of junctional mRNAs. In particular, we uncover a role for components of the EJC, in particular MAGOH, in cell junction targeting and polarity regulation, a function that is conserved evolutionarily. Moreover, we find that MAGOH and ZO-1 expression is correlated in high-grade serous epithelial ovarian cancer (HGS-EOC) tissues and that their reduced expression correlates with poor disease outcome. Altogether, this study highlights the importance of local mRNA regulation at the level of cell junction structures for epithelial cell polarity regulation.

## RESULTS

### Conserved cortical targeting and localized translation of mRNAs encoding cell junction proteins

In a previous large-scale FISH screen for localized mRNAs in *Drosophila*, several transcripts were found to localize to distinct territories along the lateral membrane of embryonic epithelial cells formed during cellularization [27]. This included the *Pyd*, *Cno* and *Scrib* transcripts **(Figure 1A)**, which encode protein components of cell junction structures required for the establishment and maintenance of epithelial cell polarity [33, 34]. We first sought to assess whether this junctional localization is observed in other epithelial tissues or cells. Comparative FISH analyses of *Drosophila* embryos and dissected ovarian egg chambers revealed the consistent targeting of the *Pyd*, *Cno* and *Scrib* mRNAs to cell junction regions along the lateral membranes embryonic and follicular epithelial cells, respectively **(Figure 1A-B)**. We next extended these analyses to the human breast carcinoma cell line MCF7, which exhibits robust cobble-stone localization of adherens (E-Cadherin/Ecad) and tight (Zonula Occludens-1/ZO-1) junction proteins after 3 days in culture. FISH on MCF7 cells revealed that *ZO-1*, *Afadin/Afdn* and *Scrib* mRNAs, respective orthologues of *Pyd*, *Cno* and *Scrib*, also display cobble stone-like cell junction patterns in confluent cellular monolayers **(Figure 1C)**. Immuno-FISH assays revealed that the junctional distribution patterns of *Zo-1* mRNA appears to precede, or coincide with, the targeting of its protein in MCF7 cells **(Figure 1D)**. These results reveal that cell junction targeting is a conserved feature of epithelial polarity-associated mRNAs.

**Figure 1.**
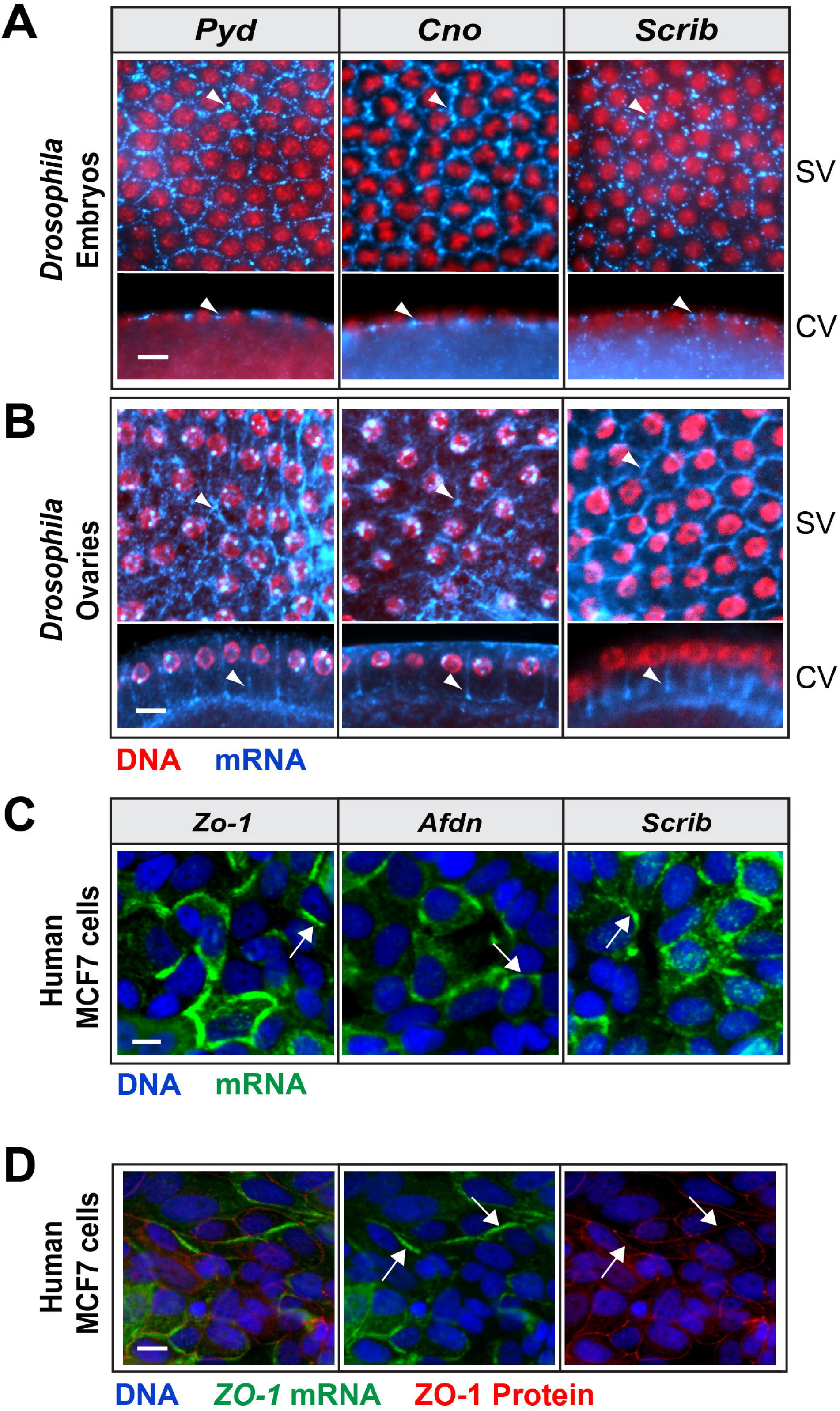
Localization of mRNAs to cell junction regions in *Drosophila* tissues and mammalian epithelial cells. **(A-B)** RNA FISH of wild-type (*OreR*) *Drosophila* embryos (A) or egg chambers (B) revealing the localization of *Pyd, Cno, and Scrib* mRNAs (blue) to the lateral membrane of embryonic epithelial cells (A) or follicular epithelial cells (B). Red, DNA. SV, surface view. CV, cross-section view. Scale bar represents 10µm for (A) and 15µm in (B). **(C)** RNA FISH in human MCF7 cells showing the localization of *Zo-1, Afdn* and *Scrib* mRNAs (Green) to regions of cell-cell contacts. Blue, DNA. Scale bar represents 15µm. **(D)** RNA FISH coupled with IF of MCF7 cells reveals that *Zo-1* mRNA (green) localization at regions of cell-cell contacts in relation to ZO-1 protein (red). RNA and protein of Zo-1 are partially co-localized at regions of cell-cell contacts (arrows). Blue, DNA. Scale bar represents 15µm.

To assess whether cell junction transcripts undergo localized translation, we next employed puromycin (Puro) labeling combined with proximity ligation assays (Puro-PLA), a recently developed method to study localized translation in cell culture or tissue specimens [35–37]. As detailed in **Figure 2A**, this approach involves brief exposure of cells to Puro to inhibit translation and cause the release of truncated puromycylated protein products, followed by rapid fixation to prevent molecules from dispersing away from their site of synthesis. Immunolabeling is then conducted with an anti-Puro antibody, to label puromycylated proteins, and an antibody recognizing a protein of interest, followed by secondary antibodies conjugated to oligonucleotide probes that allow rolling-circle signal amplification when in close proximity. The amplified products are discernable microscopically as fluorescent foci that represent sites of nascent synthesis of the targeted protein [35]. We first performed Puro-PLA analyses on MCF7 cells at different days post-seeding with a rabbit anti-SCRIB antibody, which revealed prominent nascent protein foci in the cell periphery over a 3-day time course **(Figure 2B)**. Interestingly, more Puro-PLA foci were observed at early time points, suggesting that localized translation is an early event that may contribute to polarity establishment. To compare the distribution properties of the SCRIB/Puro-PLA signal to sites of SCRIB protein accumulation, a fluorescent anti-rabbit secondary antibody was added to specimens after the Puro-PLA reactions were completed. While the signal was generally non-overlapping, due to competition between secondary antibodies for common epitopes, we observed some sites where the Puro-PLA and immunofluorescence (IF) signals were co-localized in the cell cortex. Similarly, cortical Puro-PLA patterns were observed for ZO-1 and AFDN labelled samples **(Figure 2C)**, indicating that these corresponding transcripts also undergo localized translation in the cell periphery. By contrast, the Puro-PLA signal was absent in non Puro-treated cells **(Figure 2B-C)**, in cells lacking primary antibodies, or in cells treated with anisomycin, an inhibitor of ribosomal peptidyl transferase activity, prior to Puro exposure **(Figure 2C)**. We conclude that the *Scrib*, *Zo-1* and *Afdn* mRNAs undergo localized translation at cell junction regions, sites where their encoded proteins are known to function.

**Figure 2.**
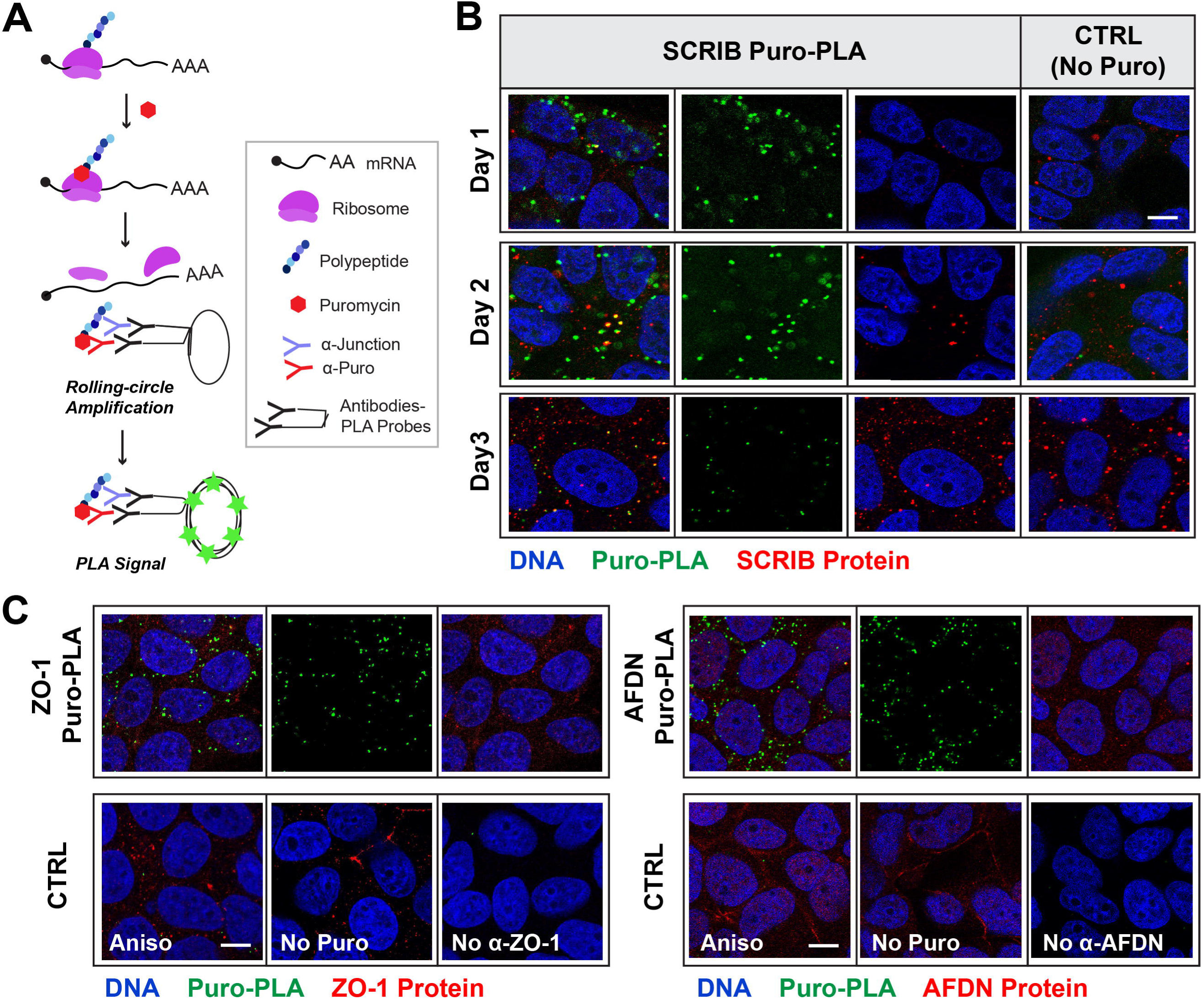
Cell Junction mRNAs are locally translated at cell junction regions. **(A)** Schematic representation of Puro-PLA assay to visualize sites of nascent translation. **(B)** Confocal imaging of SCRIB/Puro-PLA (green) coupled with SCRIB IF (red) at different days post-seeding of MCF7 cells. SCRIB exhibits localized translation in regions of cell-cell contact, as revealed by robust SCRIB/Puro-PLA signal in day 1 and day 2 cells, with a reduction by day 3. Conversely, the SCRIB protein IF signal at the cell junction regions becomes more pronounced by day 3. Lack of Puro treatment results in an absence of Puro-PLA signal, demonstrating specificity of the reaction (right panels). Blue, DNA. Scale bar represents 10µm. **(C)** Imaging of ZO-1/Puro-PLA and AFDN/Puro-PLA signals (green), contrasted to their respective protein IF (red), reveals localized translation at cell-cell contact regions (upper panels). As negative controls, samples pretreated with anisomycin, non-Puro treated, or lacking primary antibodies were generated (bottom panels). Blue, DNA. Scale bar represents 10μm.

### Imaging screens identify a variety of cell junction localized RBPs

The trafficking and localized translation of mRNAs is typically coordinated by *trans-*acting RBPs. To identify potential regulators of cell junction transcripts, we first conducted a high-content imaging screen in polarized MCF7 cells **(Figure 3A)**, using a collection of ∼400 antibodies directed against proteins bearing canonical RNA binding domains (RBDs) or identified in RNA interactome studies [38, 39]. These RBP antibodies were thoroughly validated for specificity via immunoprecipitation (IP), Western-blotting and loss-of-function (LOF) methodologies [40], and were recently used to study the subcellular distribution patterns of RBPs by systematic IF labeling under steady-state or stress conditions in different cellular models [41, 42]. Collectively, the interrogated proteins have been implicated in a variety of post-transcriptional regulatory steps (e.g. RNA splicing, stability, translation), although many have yet poorly defined functions in RNA metabolism.

**Fig. 3.**
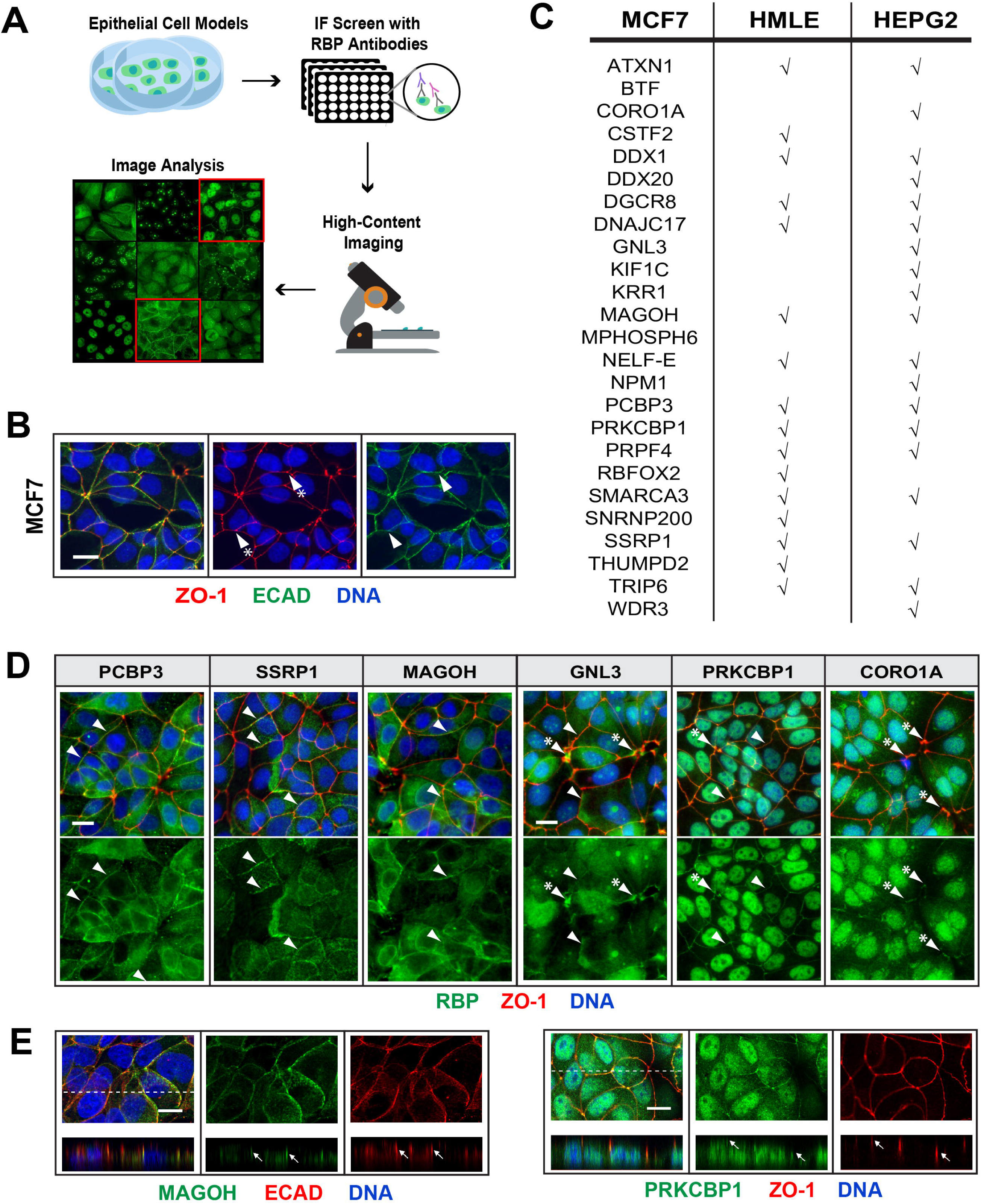
High content imaging screens for cell junction localized RBPs. **(A)** Schematic outline of high-content imaging IF-based screens for RBPs that localize to cell junction regions in MCF7, HepG2 and HMLE cells. **(B)** Representative IF images of ZO-1 (red) and E-CAD (green) co-labeling in MCF7 cells at Day 3 post-seeding. Arrowhead with star points to tri-cellular enrichment in ZO-1. Blue, DNA. Scale bar represents 30µm. **(C)** RBPs with junctional distribution patterns detected by IF in MCF7 cells and results of validation assays in HMLE and HEPG2 cells. **(D)** Representative images for candidate RBPs (green) with cell junction IF patterns in relation to ZO-1 protein (red). Some RBPs are broadly distributed through the junctional regions (e.g. PCBP3, SSRP1, MAGOH), while others appear more enriched at tri-cellular junctions (e.g. GNL3, PRKCBP1, CORO1A). Arrowheads illustrate areas of co-localization with ZO-1, while arrowheads with stars indicate pronounced tri-cellular IF signals. Blue, DNA. Scale bar represents 30µm. **(E)** Confocal z-stack renderings were generated at 0.5μM intervals to further validate the co-localization of cell junction RBP candidates with cell junction markers, as shown by the arrows. As examples, MAGOH co-labeling with E-CAD (left panel) and PRKCBP1 co-detection with ZO-1 (right panels) are shown. Arrows indicate sites of signal overlap. Dotted lines indicate the sites from which z-stacks are derived. Green, MAGOH and PRKCBP1. Red, E-CAD and ZO-1. Blue, DNA. Scale bar represents 15μm.

MCF7 cells were cultured for 3 days post-seeding on 96-well plates and were subjected to systematic IF to co-visualize each candidate RBP in relation to the ZO-1 protein, identifying 25 RBPs with cell junction-like patterns **(Figure 3B-C; Table S1)**. This included proteins such as PCBP3, SSRP1 and MAGOH, which showed a broad distribution along the length of the cell-cell contact surfaces, as well as factors such as GNL3, PRKCBP1 and CORO1A that tended to be more enriched at tri-cellular junction sites **(Figure 3D)**. Through confocal imaging and z-stack renderings, we further validated the cell junction targeting behavior of these RBPs, as shown for MAGOH and PRKCBP1, which respectively co-localized with canonical cell junction makers, E-CAD and ZO-1 **(Figure 3E)**. We next tested whether the cell junction targeting behaviour of these proteins was manifest in other epithelial cell models by performing comparative IFs in non-tumorigenic human mammary epithelial (HMLE) cells and in the HEPG2 liver hepatocarcinoma cell line. The demarcation of cell junction regions with the ZO-1 marker was less robust in these cells compared to MCF7, requiring prolonged incubation times **(Figure S1)**. Nevertheless, many of the RBP candidates identified in MCF7 cells were also found to be junctional in HMLE and HEPG2 cells (**Figure 3C**). These RBP hits include factors involved in mRNA processing (e.g. RBFOX2, PRPF4, SNRNP200), RNA helicases (e.g. DDX1, DDX20) and proteins implicated in non-coding RNA biogenesis (e.g. NPM1). Consistent with previous observations that microRNA regulatory factors are found at adherens junctions [32], we observed junctional-targeting of DGCR8, a component of the microprocessor complex implicated in pre-miRNA maturation. Interestingly, several proteins of the EJC, including MAGOH, RBM8A/Y14 and EIF4A3, were found to partially localize to cell junction regions **(Figure S2)**. Altogether, these results reveal that a substantial fraction of interrogated RBPs (∼6%) localize to cell junction regions in epithelial cells, potentially implicating these factors in localized RNA regulation at these structures.

### Candidate RBPs physically associate with cell junction mRNAs

To investigate the mRNA target repertoire of RBPs that localize to cell junctions, we first interrogated recently published enhanced cross-linking and IP (eCLIP) data [42], an assay that measures the binding properties of a given RBP in a transcriptome-wide fashion. Using available eCLIP datasets for 5 cell junction RBPs (BTF, CSTF2, PRPF4, RBFOX2 and SMARCA3), as well as 3 control RBPs that are not cell junction localized (PABPN1, SFQPQ and U2AF1), we evaluated the functional signatures of their target mRNAs via gene ontology (GO) term enrichment analyses, focusing on cell component GO terms with adjusted p-values < 0.05 **(Figure 4A)**. These analyses revealed commonly enriched GO terms across all RBPs, including chromosome, nucleolus, and ribonucleoprotein complex, while the cell junction RBPs exhibited more selective enrichments for terms related to nuclear matrix, adherens junctions and apical plasma membrane. We also took advantage of the CLIPdb database [43], an online resource with curated binding site information for hundreds of published datasets generated using different CLIP-seq methodologies (e.g. HITS-CLIP, iCLIP, PAR-CLIP, eCLIP), to assess whether direct binding associations for specific RBPs have been mapped to various cell junction mRNAs. As shown in **Figure 4B**, of the eight junctional RBPs with curated datasets in CLIPdb, which were generated using HEPG2 or HEK293 cells, a broad set of direct interactions are observed between these factors and cell junction mRNAs.

**Fig. 4.**
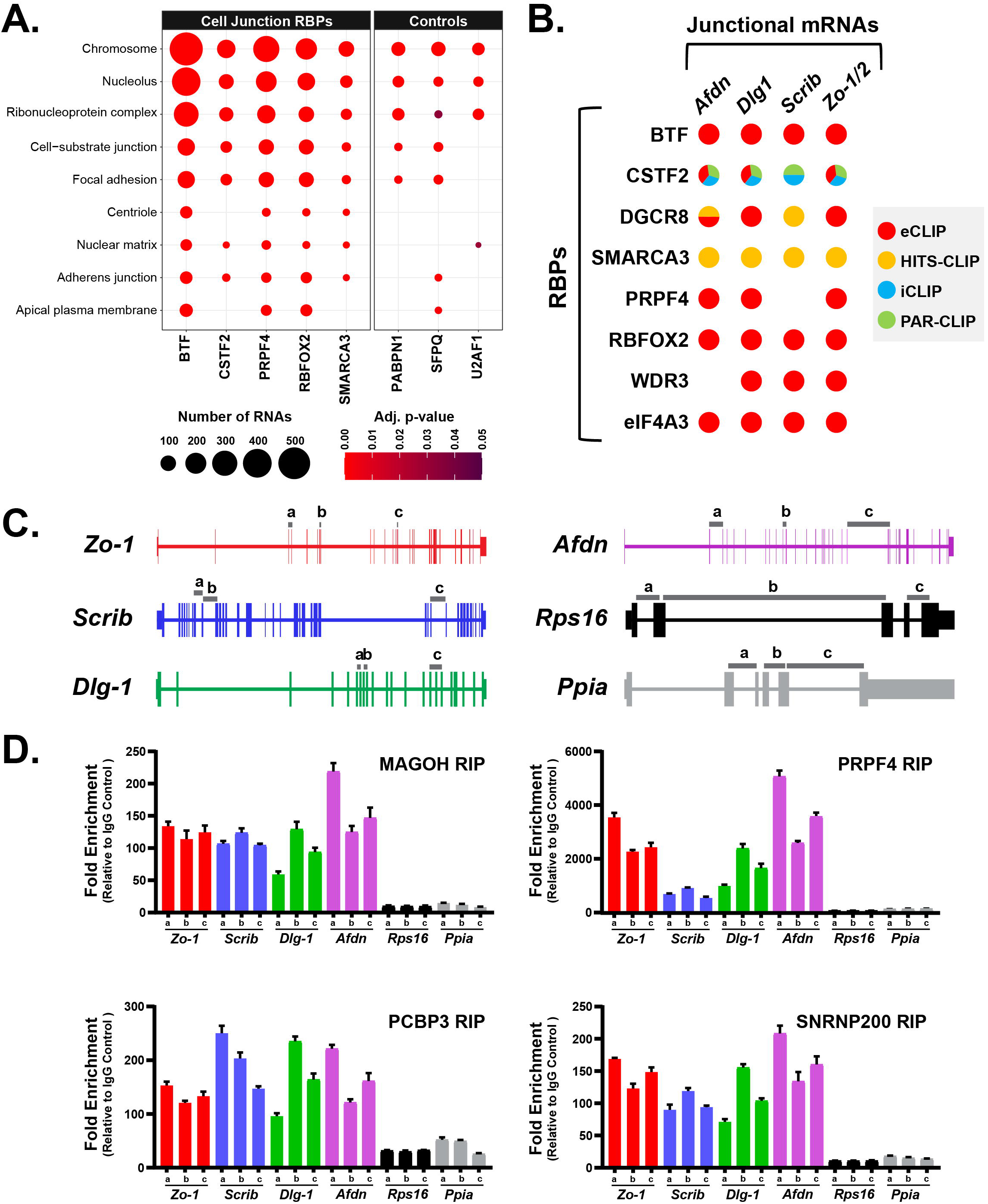
Junctional RBPs exhibit robust physical association with cell junction mRNAs. **(A)** Bubble plots of cell component GO-term enrichment analyses of public eCLIP data for the indicated RBPs [42], including 5 junctional RBP candidates and 3 non-junctional factors. The number of bound mRNAs associated with specific GO categories is represented as a function of circle size, while the red gradient scale represents adjusted p-values. **(B)** Survey of CLIPdb datasets to identify binding contacts of cell junction RBP candidates to mRNAs that localize to cell junction regions, assessed using different CLIP procedures. **(C)** Schematic representation of transcript-specific tiling primer sets used for detect expression of junctional (*Zo-1, Scrib, Dlg1, Afdn*) and control (*Rps16, Ppia*) mRNA candidates in RIP/RT-qPCR assays conducted in (D). **(D)** RIP/RT-qPCR assays performed with MCF7 cellular extracts using antibodies for the indicated junctional RBP candidates or control IgG. RNA recovered from each immunoprecipitation was extracted and processed by RT-qPCR using the indicated (*a, b, c*) primer pairs represented in (C) for each detected mRNA. Fold enrichments were calculated relative to IgG control samples after normalization to their respective inputs. Data are shown as mean ± SD, n=3.

To validate the physical associations of cell junction localized RBPs with junctional mRNAs in MCF7 cells, we next performed RNA immunoprecipitation (RIP) of endogenous candidate RBPs, including MAGOH, PCBP3, PRPF4 and SNRNP200, using whole cell lysates, followed by quantitative reverse transcription PCR (RT-qPCR) analysis. For each analyzed transcript, which included junction localized (*Zo-1, Scrib, Dlg-1* and *Afdn*) and non-junctional controls (*Rsp16, Ppia*), we designed oligonucleotide pairs probing different regions of the mRNA sequence to verify signal consistency **(Figure 4C)**. Notably, all four interrogated RBPs exhibited strong enrichments, relative to control purifications conducted with non-specific IgG, for junctional mRNAs when compared to non-junctional transcripts **(Figure 4D)**. While the fold enrichment values relative to IgG control samples varied depending on the RBP, the co-IP efficiency was systematically higher for junctional transcripts compared to control mRNAs and the signals were consistent across oligonucleotide pairs used for each mRNA. Altogether, these results reveal that several cell junction RBPs exhibit preferential physical association with junction-localized mRNAs.

### Functional disruption of junctional RBPs perturbs cell junction marker patterning

We next sought to assess the impact of functional disruption of cell junction RBPs on epithelial cell organization. For this, we conducted siRNA-mediated depletion of cell junction RBPs, as well as control scrambled siRNA, and quantified the impact of these perturbations on the targeting properties of ZO-1 and SCRIB along the lateral membranes of MCF7 cells (**Figure 5A**). These proteins label slightly overlapping territories along the lateral membrane, with ZO-1 having a tighter apical localization and SCRIB showing a broader basolateral distribution. Therefore, we analyzed 3D confocal z-stack renderings of MCF7 cells treated with scrambled or RBP-specific siRNAs, in order to define the impact of RBP depletion on the thickness (in µm) of the lateral membranes, as well as the centroid positioning (expressed as a percentage of cell length) of SCRIB and ZO-1 localization territories **(Figure 5A-C)**. The tested siRNAs were generally found to efficiently deplete targeted RBP mRNA levels by > 70%, as assessed by RT-qPCR analysis **(Figure 5B, upper row of heatmap)**. Strikingly, knock-down of most cell junction RBP candidates altered the thickness of MCF7 cell lateral membranes, as well as the thickness of the SCRIB domain and/or ZO-1 localization (**Figure 5B**). For instance, depletion of PCBP3, MAGOH and SSRP1 let to a weakening or disruption of the normally tight and organized junctional distribution properties of SCRIB and ZO-1 **(Figure 5C)**, phenotypes that were quantified across a minimum of 50 cell-cell borders per knock-down condition **(Figure 5D-E)**. We conclude that the functional disruption of many cell junction RBPs leads to perturbations in the normal distribution of junctional proteins encoded by localized mRNAs.

**Fig. 5.**
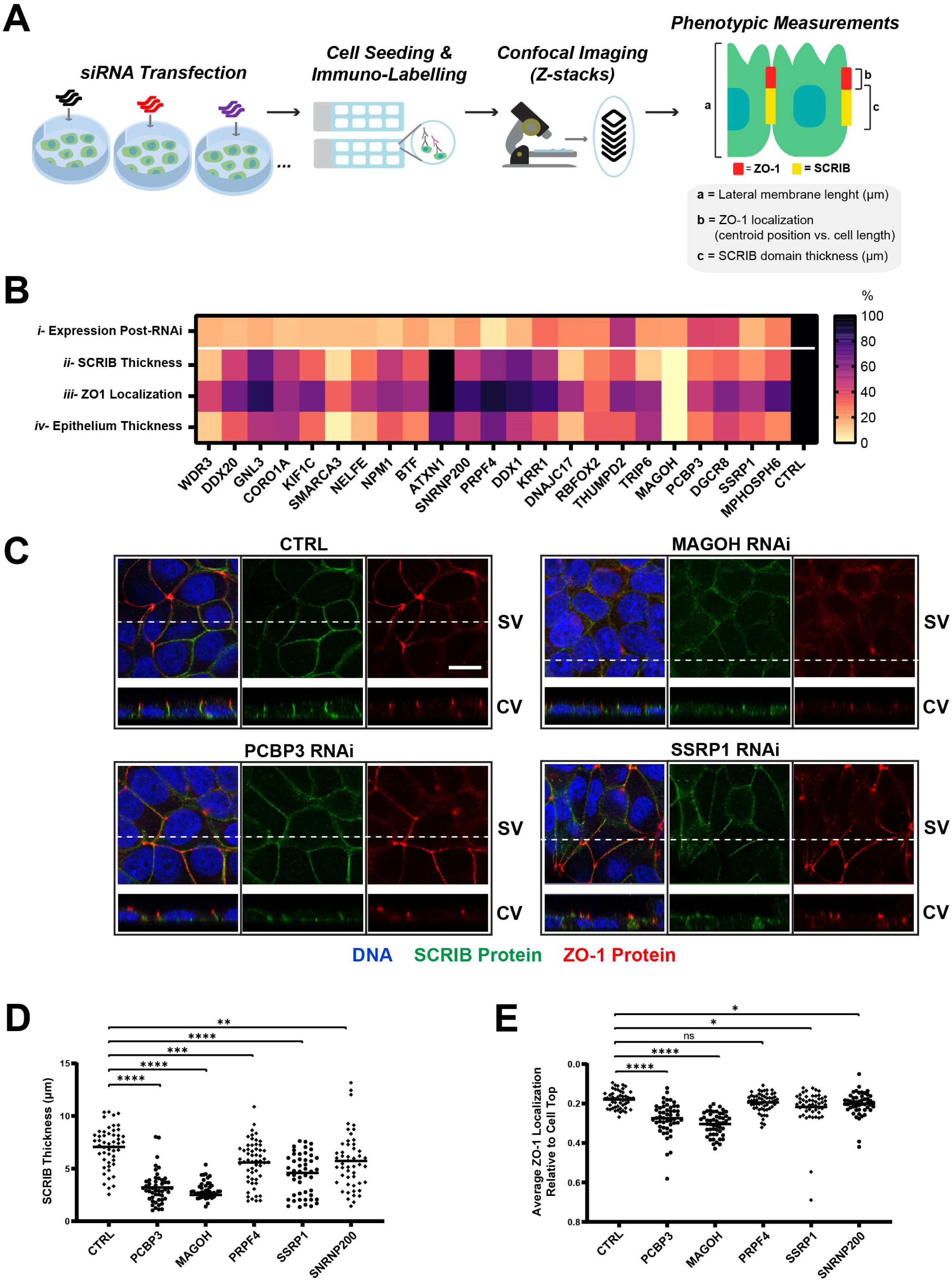
Impact of RBP loss-of-function on epithelial cell polarity and cell junction marker distribution. **(A)** Schematic outline of siRNA transfection approach to deplete candidate RBPs in MCF7 cells. Following siRNA transfections, cells were seeded for 3 days and processed for IF to visualize and quantify ZO-1 and SCRIB protein distribution phenotypes in confocal Z-stack renderings. **(B)** Heat map depicting quantitative phenotypic read outs in MCF7 cells following RBP-RNAi treatments, represented as percentage relative to scrambled-RNAi controls. Quantifications include (i) the average percent normalized expression values of candidate RBP transcripts following RNAi, (ii) the average percent thickness of SCRIB localization domain, (iii) the average percent localization of ZO-1 relative to cell top, and (iv) the average percent thickness of the epithelium. **(C)** Representative examples of IF results for ZO-1 (red) and SCRIB (green) proteins in control versus RBP-RNAi in MCF7 cells. Typically, SCRIB localizes broadly along the lateral membrane, while ZO-1 occupies a more discrete apical territory, as revealed by confocal microscopy image Z-stacks taken at 0.5μM intervals. Note that siRNA-mediated knockdowns of candidate RBPs, including PCBP3, MAGOH and SSRP1, led to changes in SCRIB thickness and ZO-1 localization along the lateral membrane, compared to control. Dotted lines in the surface view (SV) images mark the sites at which the cross-section views (CV) were generated. Blue, DNA. Scale bar represents 15 μm. **(D)** Quantification of the impact of RNAi treatments on SCRIB domain thickness (µm) following treatment with Control, PCBP3, MAGOH, PRPF4, SSRP1 and SNRNP200 siRNAs. Statistical significance was assessed by two-tailed t-tests; ****, *** and ** denote p-value of <0.0001, 0.001 and 0.01, respectively. **(E)** Quantifications of the impact of RNAi treatments with Control, PCBP3, MAGOG, PRPF4, SSRP1 and SNRNP200 siRNAs on the average localization of ZO-1 relative to the top of the cell on a scale of 0 to 1. Statistical significance was assessed by two-tailed t-tests; **** and * denote p-value of <0.0001 and <0.05, respectively, while ‘ns’ denotes non-significance.

### MAGOH and PCBP3 are required for localized translation of junctional mRNAs

We next investigated whether candidate RBPs important for proper cell junction protein patterning are also required for localized translation of the mRNAs encoding these proteins. To address this question, Puro-PLA for SCRIB or ZO-1 was performed on MCF7 cells treated with siRNA targeting MAGOH, PCBP3 or scrambled control, and the samples were then imaged to generate Z-stack renderings. After completion of the Puro-PLA reactions, IF detection of residual SCRIB and ZO-1 protein signal was performed in order to demarcate cell junction regions for downstream image analysis. As shown in **Figure 6A**, in control scrambled siRNA samples, Puro-PLA for SCRIB and ZO-1 revealed an abundance of translation foci within the cell junction territories. By contrast, treatments with siRNAs targeting MAGOH and PCBP3 strikingly reduced the number of Puro-PLA foci at cell-cell contacts compared to control cells. To quantify junctional Puro-PLA foci more precisely, masks covering the cell junction regions of Z-stack reconstructions were created and junction-proximal Puro-PLA foci were quantified **(Figure 6B-C)**. To confirm the specificity of the assay, control reactions were performed in which Puro treatment was omitted or that were pre-treated with anisomycin prior to Puro exposure, revealing a near complete loss of Puro-PLA foci **(Figure 6A, C)**. Together, these results indicate that MAGOH and PCBP3 are required for the localized translation of *Scrib* and *Zo-*1 mRNAs at cell junction regions.

**Fig. 6.**
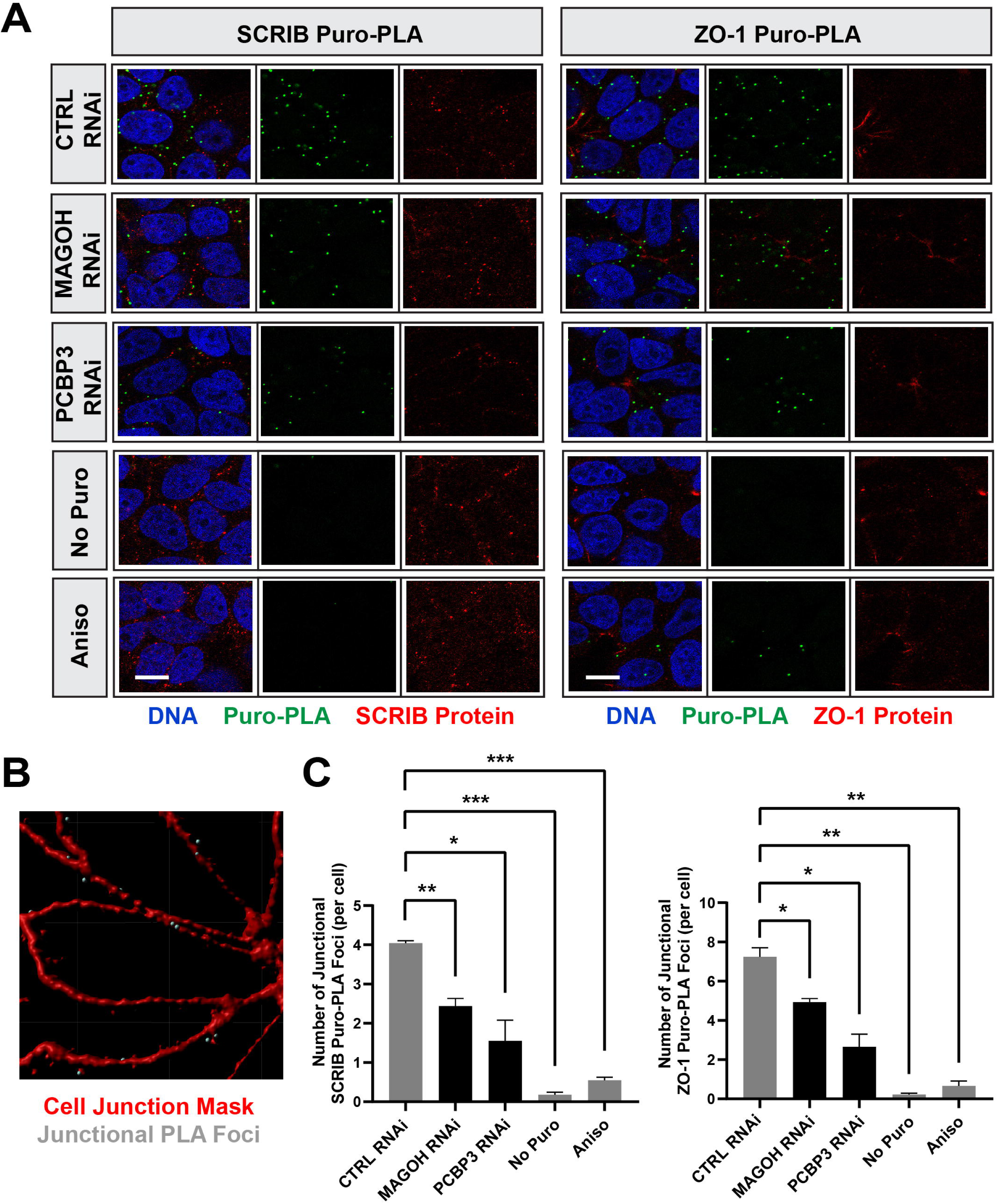
Assessing the impact of RBP-depletion on localized translation of polarity and cell junction factors. **(A)** SCRIB/Puro-PLA and ZO-1/Puro-PLA assays were performed on MCF-7 cells following treatment with Control, PCBP3 or MAGOH siRNAs. Note that knockdown of MAGOH or PCBP3 reduces localized translation of both SCRIB and ZO-1, as witnessed by the reduced Puro-PLA signal. Cells processed in the absence of Puro treatment (No Puro), or that were treated with Anisomycin (Aniso) prior to Puro exposure, were included as controls for Puro-PLA signal specificity. Green, Puro-PLA. Red, SCRIB or ZO-1 protein. Blue, DNA. Scale bar represents 10µm. **(B)** Representative image of cell junction mask (red) used for quantifying Puro-PLA foci (grey) at cell membrane regions using Imaris software. **(C)** Quantification of the number of Puro-PLA foci demonstrating co-localization cell junction masks on 3D confocal z-stack renderings. Statistical significance was assessed via two-tailed t- tests; ***, ** and * denote p-value of <0.001, 0.01 and 0.05, respectively. Error bar denotes SD, n=60 or more.

### Disruption of EJC components perturbs junctional mRNA and protein distribution in *Drosophila*

To assess whether EJC components influence epithelial cell polarity in an evolutionarily conserved fashion, we next utilized the Gal4-UAS system in *Drosophila* to perturb expression of Mago Nashi (MAGO), the fly orthologue of MAGOH, *in vivo* in follicular epithelial cells. For this, expression of a hairpin RNA (hpRNA) targeting *mago* mRNA, under control of an upstream activating sequence (UAS) element, was induced in the follicular epithelium using the *traffic jam-Gal4* (*TJ-Gal4*) driver. Strikingly, in comparison to control specimens, *mago-RNAi* altered the junctional distribution of both PYD protein and *Pyd* mRNA in late-stage egg chambers, while also perturbing the integrity of the epithelial cells layer (**Figure 7A-B**). We also assessed the impact of EJC component depletion on the distribution of DLG1 protein, a component of the Scribble polarity complex for which the mRNA is also known to localize to cell junction regions in *Drosophila* [27]. Stikingly, *mago-RNAi* egg chambers exhibit disrupted morphology and a strong reduction in DLG1 protein distribution in follicular epithelial cells (**Figure 7C**). Since MAGO is one of four core components of the splicing-dependent EJC, which also includes the Tsunagi/TSU, EIF4A3 and Barentsz/BTZ proteins, we subsequently investigated whether loss-of-function of these EJC components produces similar phenotypes. Strikingly, *tsu-RNAi* phenocopied *mago-RNAi*, including perturbations in polarity marker distribution and misalignment of the nuclei along the follicular epithelium. By contrast, RNAi-mediated depletion of *eif4a3* proved lethal, while *btz-RNAi* gave no phenotype (not shown). We conclude that MAGO and its dimerization partner TSU are required for the proper targeting of junctional proteins/mRNAs and for the maintaining epithelial polarity in follicular epithelial cells.

**Fig. 7.**
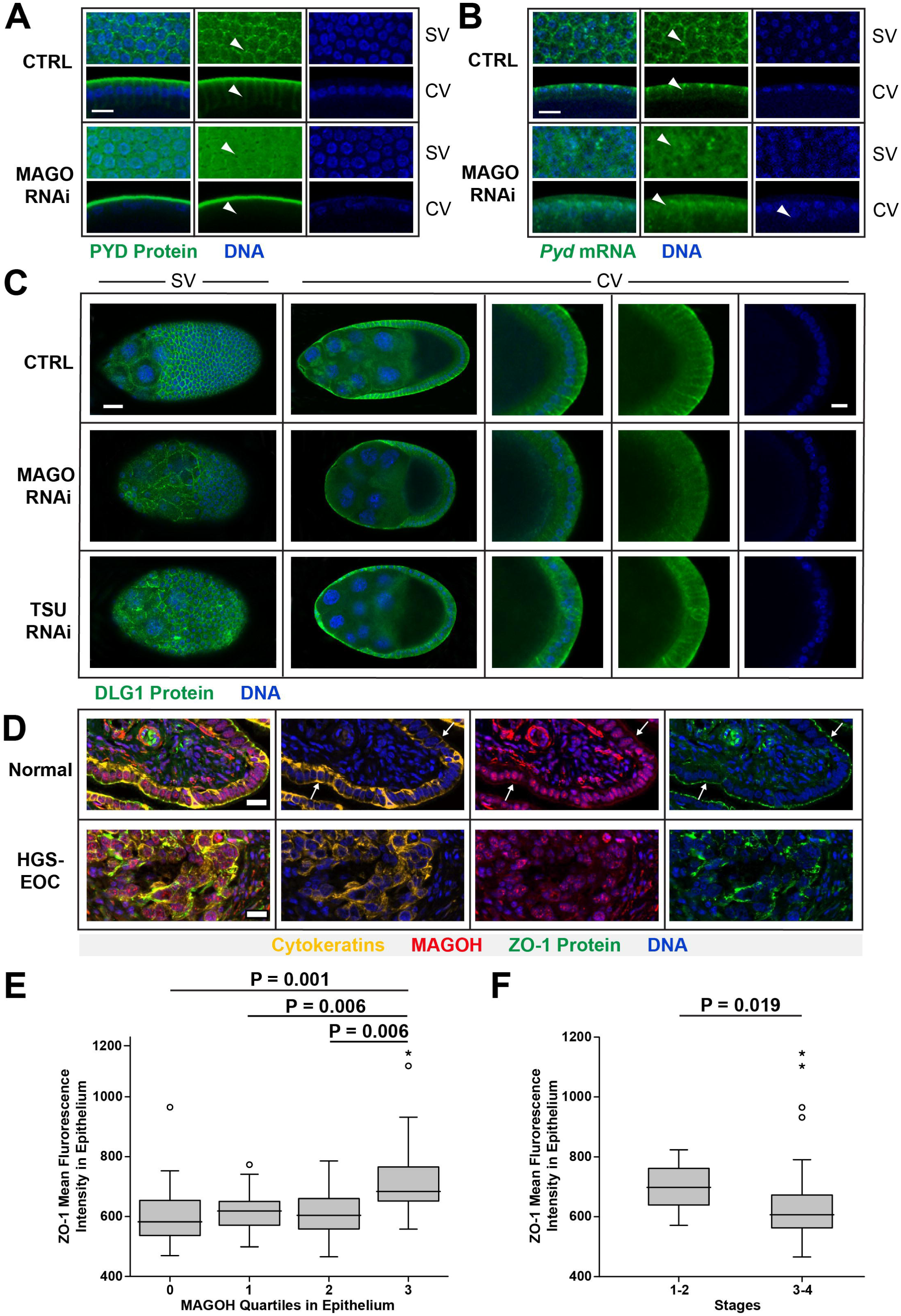
***In vivo* loss-of-function studies for PYD and DLG1 in *Drosophila* egg chambers as well as surgical characteristics of MAGOH and ZO-1 in human ovarian tissues. (A-B)** Visualization of PYD protein by IF (A) and *pyd* mRNA by FISH (B) in follicular epithelial cells of control or *MAGO-RNAi* egg chambers of female *Drosophila* resulting from crosses of *Tj-Gal4* females to *UAS-MAGO-hpRNA* male flies. Egg chambers of *TJ-Gal4* females were used as controls. Green, PYD protein (A) or *pyd* mRNA (B). Blue, DNA. SV, surface view. CV, cross-section view. Scale bars represent 15µm. **(C)** Visualization of DLG-1 protein (green) distribution by IF in control (*TJ-Gal4*), *MAGO-RNAi* or *TSU-RNAi* egg chambers, resulting from crosses as in (A-B). The first column represents the surface views (SV), while other columns are the cross-section view (CV). Squares represent the enlarged view of the epithelium. Blue, DNA. Scale bar on the left and right represent 15µm and 5µm, respectively. **(D)** Representative TMA staining of Cytokeratins (yellow), MAGOH (red) and ZO-1 (green) proteins in a discovery cohort composed of control human fallopian tube tissue specimens (n = 15) and high-grade serous epithelial ovarian cancer (HGS-EOC) specimens (n = 101). Arrows point to MAGOH staining at the apical pole of the epithelial cells in normal tissue, illustrating partial co-localization with cytokeratin and ZO-1 markers. Blue, DNA. Scale bars represent 20µm. **(E)** Box plot illustrating the mean fluorescence intensity (MFI) of ZO-1 protein in epithelia of HCS tissues (n = 90) in relation to MAGOH protein MFI, which was separated into 4 quartiles ranging from negative (0, n = 22), low (1, n = 22), moderate (2, n =23) and high (3, n =22), using approximately the 25^th^, 50^th^ and 75^th^ percentiles as cut-offs between these groups. Notably, high MAGOH expression in epithelial structures is positively correlated with heightened ZO-1 expression, as seen by the significant distinction between group 3, denoting high expression, and all other groups (p = 0.001, 0.006, 0.006, when compared to group 0, 1, 2, respectively, as assessed via 2-tailed T-tests). Both protein MFI levels were digitally quantified in an automated fashion. Boxes equal the interquartile (IQ) range of samples, while whiskers represent the highest and lowest values, which are less than 1.5 times the range of IQ. Circles denote the outliers and stars highlight the extreme outliers, which deviate from the IQ range by more than 1.5 times or 2 times the IQ range, respectively. **(F)** Box plot showing the MFI distribution of ZO-1 in early (1-2) vs. late tumor (3-4) stages, characterized according to international standards. A significant reduction of ZO-1 expression is noted in epithelial tissues of later or more severe stages of HGS-EOC (n = 80) compared to early stages (n = 12). The smaller sample size in early tumor stages is due to the low frequency of women being diagnosed with stage 1-2 cancer. The two-tailed t-test was utilized to compare the means, where p = 0.019.

### Evaluating MAGOH and ZO-1 expression features in ovarian tumor specimens

We next sought to examine the expression properties of ZO-1 and MAGOH in the context of control and ovarian cancer tissue specimens by conducting IF co-labeling studies on tissue microarrays (TMAs). For this, we utilized arrays comprising a discovery cohort of 15 normal fallopian tube specimens and a tumor panel of 101 high-grade serous (HGS) carcinomas of uterine adnexa [44, 45], the most common and aggressive form of epithelial ovarian cancer (EOC)[46]. TMA slides were subjected to IF with antibodies for ZO-1, MAGOH and epithelial cytokeratins (**Figure 7D and S3A**), and were then subjected to computerized image analysis to detect regions of interest, such as epithelial structures and stroma. The mean fluorescence intensity (MFI) of MAGOH and ZO-1 protein expression within the epithelial compartment was measured and these values correlated to associated clinical data in order to evaluate potential links to disease prognosis. Consistent with existing literature, ZO-1 robustly localizes to the apical domain of epithelial cells in normal fallopian tubes (**Figure 7D**, top panels) [47]. However, the apical distribution of ZO-1 was perturbed in epithelial tissue of HGS specimens, depicting a loss of apico-basal polarity (**Figure 7D**, bottom panels). While MAGOH exhibited a stronger enrichment within the epithelial cell nuclei and surrounding stroma, its targeting to the apical pole of epithelial cells was observed in normal tissues, a feature that is diminished in HGS tissues **(Figure 7D)**. Cytokeratin staining revealed a monolayer of nuclei in epithelia of normal fallopian tube specimens, which was largely altered in HGS-EOC, further supporting a loss of epithelial cell polarity. Comparative analyses revealed that high MAGOH expression in epithelial structures is positively correlated with heightened ZO-1 expression (**Figure 7E**), as seen by the significant distinction between group 3 and lower MAGOH expression quartiles [Q0-2] (p = 0.001-0.006), as well as a highly significant positive Spearman correlation between MAGOH and ZO-1 protein levels (Rho = 0.430, P < 0.001, n = 89) in HGS-EOC tissue. Interestingly, analysis of ZO-1 expression in relation to disease progression stages revealed a significant tendency (p = 0.019) for lower expression in the more advanced tumors (stages 3-4) compared to early stage 1-2 specimens (**Figure 7E**). Moreover, correlating MAGOH MFI data to disease free survival and survival time reveals that a higher level of MAGOH epithelial expression shows a tendency to associate with better outcome (**Figure S3B** and **Figure S3C**), although further testing on TMAs of larger cohort size would be required to establish more robust significance. Taken together, these observations suggest that ZO-1 and MAGOH expression is correlated in HGS-EOC, consistent with the regulatory links described above, while also suggesting that they may represent potential biomarkers of good prognosis in ovarian cancer.

## DISCUSSION

The polarization of epithelial cells along their apico-basal axis is a highly conserved feature in metazoan organisms, which is imparted by the demarcation of distinct membrane territories and cell junction structures required for tissue integrity and the selective passage of nutrients from the external environment [2]. The establishment and maintenance of epithelial polarity has traditionally been attributed to the asymmetric distribution of specific lipid species and protein modules [1, 6]. However, a significant body of literature has also revealed that RNA molecules can be selectively partitioned within different cytoplasmic compartments and membrane structures of epithelial cells [23, 27, 28, 30, 32, 48–52], although the impact of post-transcriptional regulatory events on epithelial cell polarization remains unclear. In the present study, through comparative analyses in *Drosophila* and human epithelial tissues and cell models, we show that the junctional targeting of several mRNAs encoding cell junction and polarity modulating proteins is a conserved feature of epithelia across evolution. Our finding that the localization of these mRNAs typically precedes robust cell junctional targeting of their encoded proteins suggested that they may undergo localized translation, a hypothesis that we confirm using Puro-PLA to label sites of nascent protein synthesis for ZO-1, AFDN and SCRIB. Through high-content imaging with a validated antibody collection targeting ∼400 human RBPs, we further identify a collection of RBPs demonstrating steady-state localization to cell junction regions, several of which directly physically interact with junction targeted mRNAs. RNAi-mediated functional disruption of these RBPs revealed their requirement for proper partitioning of epithelial polarity marker proteins. Focusing on two such candidates, PCBP3 and MAGOH, which have previously been linked to diverse RNA regulatory functionalities, we find that their depletion disrupts the localized translation of *Scrib* and *Zo-1* mRNAs at cell junction regions. Finally, we found that *in vivo* functional disruption of *Drosophila* MAGO (i.e. the fly orthologue of MAGOH) in follicular epithelial cells, as well as other components of the EJC, perturbs the localization of junctional mRNA/proteins and disrupts tissue morphology. This work reveals that local modulation of mRNA translational at cell junction regions is important for epithelial polarity regulation.

### Implications of localized mRNA translation in epithelial polarity control

In recent years, the asymmetric subcellular distribution of RNA molecules has emerged as a highly prevalent process in different cell types and organisms [27–30, 53–55], with transcript distribution often displaying striking coherence with the known localization patterns and functions of the encoded proteins [27, 29, 56, 57]. Such findings have underscored the potential importance of mRNA targeting and local translation for a variety of biological processes including cell migration, cell polarity control, axonal pathfinding, synaptic plasticity, and mitotic cell division [22, 36, 58–60]. A large number of transcripts have also been shown to exhibit functionally important asymmetric localization patterns in epithelial tissues [23–25, 27, 28, 30, 32, 48–52, 61, 62]. Indeed, systematic surveys utilizing high-throughput imaging approaches or tissue micro-dissection coupled with RNA sequencing have identified a diverse repertoire of mRNAs that partition selectively along the apico-basal axis or at cell junction regions in *Drosophila*, *C elegans* and mouse epithelial tissues [27, 28, 30, 48]. Indeed, Moor et *al.* recently observed that transcripts encoding ribosomal proteins are enriched in the apical cytoplasm of mouse intestinal epithelial cells, a phenotype enhanced under feeding conditions, thus imparting a higher rate of translation in the apical cytoplasm to promote nutrient absorption [28]. In fly ovarian and embryonic epithelial cells, the apical localization of transcripts such as *Wg*, *Crb* and *Sdt* is important for proper protein targeting, epithelial polarity control and morphogen-mediated signaling events [23–25, 51, 63]. The apical targeting of several pair-rule transcripts in fly embryos, which has classically been proposed to restrict the activity of critical regulators of embryonic patterning events [50], was later found to depend on 3’UTR stem loop elements, as well as a regulatory module formed by the EGL, the adaptor protein BIC-D and the minus-end directed microtubule motor protein Dynein [64–67]. Additional studies have begun exploring the mechanism of mRNA targeting to cell junction structures, such as the localization of *β-actin* mRNA to adherens junctions structures in MDCK cells [68], which was shown to be dependent on 3’UTR zipcode elements, as well as the case involving CPEB-mediated regulation of *Zo-1* mRNA targeting in mouse mammary epithelial cells [26]. Combined with the current study, these various examples illustrate the diversity of mRNA localization events that occur within different compartments of epithelial cells, profoundly influencing the organization, polarity, and function of these cells across deep evolutionary distances. Our findings suggest that localized mRNA translation is important for modulating the precise targeting of key cell junction proteins and epithelial polarity regulators. Since some of these locally translated proteins, including ZO-1 and AFDN, are known to physically associate within localized protein complexes [69–71], it is tempting to speculate that such polarity regulating protein modules may assemble co-translationally, a phenomenon that appears to be highly prevalent in eukaryotes [72].

### The many faces of RNA regulatory events at cell junction structures

Our systematic imaging-based screen for RBPs exhibiting steady-state localization to cell junction regions identified approximately 25 proteins with diverse functions in RNA regulation, including factors involved in non-coding RNA biogenesis (e.g. NPM1, KRR1, WDR3, GNL3, DGCR8), pre-mRNA splicing and maturation (e.g. RBFOX2, PRPF4, SNRNP200, CSTF2), RNA degradation (MPHOSPH6, DGCR8), RNA helicases (e.g. DDX1, DDX20), EJC components (e.g. MAGOH), and transcription/chromatin regulatory factors (NELF-E, SMARCA3). The list also includes several enigmatic RBPs with less established roles in RNA biology (e.g. SSRP1, CORO1A, KIF1C, TRIP6, BTF, ATXN-1, PRKCBP1, SSRP1, DNAJC17, THUMPD2), but were found to co-purify with polyadenylated RNA in interactome capture studies [38, 39, 73]. Such a diverse grouping of putative RPB candidates potentially involved in targeted RNA regulation at cell junction regions is intriguing in light of the view that RNA localization is often coordinately regulated with other aspects of RNA function, such as translational and stability control [74]. These findings are consistent with previous work by Kourtidis and colleagues [31, 32], who revealed that the PLEKHA7 adherens junction protein serves as an interface for the recruitment of components of the RNA induced silencing complex (RISC) and a subset of microRNAs to cell junction regions, where they are thought to suppress expression of mRNAs such as *Jun*, *Myc* and *Sox2*. Our work confirms the targeting of DGCR8 to junctional structures, although it remains unclear whether the cell junction transcripts that we identified are subject to localized microRNA-mediated regulation. Elucidating how the diverse activities of cell junction-localized RBP machineries participate, either through parallel, collaborative or antagonistic mechanisms, to control post-transcriptional regulatory events at these sites will be a fascinating to explore. Moreover, an important next step will also be to utilize live-cell imaging strategies to study the dynamics of cell junction mRNA localization and translational activity [75–78], as well as the relationship of translational activity to epithelial cell polarity phenotypes.

We chose to focus our follow-up interrogations on two RBP candidates that emerged from our imaging-based screen, PCBP3 and MAGOH. PCBP3 is a member of a small family of KH domain proteins implicated in the control of mRNA stability and translation [79]. Our finding that PCBP3 depletion perturbs the localized translation and targeting of junctional proteins is consistent with the suspected functions of members of this RBP family. Intriguingly, PCBP3 expression was recently identified as a prognostic biomarker for improved outcome in patients with pancreatic adenocarcinoma [80] and one might suspect that its functional role in epithelial cell polarity regulation could underlie putative tumor suppressing activities of this RBP. MAGOH is a core component of the EJC, a multi-subunit complex deposited upstream of exon-exon junctions during pre-mRNA splicing, which has been implicated in modulating the nuclear export, cytoplasmic localization, translation and stability of its bound mRNAs [81–83]. The EJC is composed of a trimeric core of proteins, including the eIF4A3 helicase that directly binds RNA and gets locked into place through interaction with a dimer formed by MAGOH and the RBM8A/Y14 protein [81, 84, 85]. The EJC core can interact with a variety of accessory factors in a cell-compartment-specific manner [82, 83, 85] and, since EJC complexes are evicted from the mRNA by ribosomes during the first pioneer round of translation, their association with a given mRNA implies that the transcript has not yet been translated. The EJC also plays a key role in the process of nonsense-mediated decay (NMD), whereby mRNAs bearing premature stop codons or containing an exon-exon junction in their 3’UTR (designated as natural NMD targets), on which at least one EJC will remain associated with the mRNA following translation, will be targeted for degradation through EJC-mediated recruitment of the NMD degradation machinery [83]. Previous studies have shown that EJC components are required for the localization of specific mRNAs, such as the targeting of *Oskar* transcripts to the posterior region of *Drosophila* oocytes [86–88]. Kwon et al. also recently showed that core EJC components are required to localize select mRNAs to basal bodies in mouse neural stem cells and human retinal pigment epithelial cells, with EJC depletion leading to defects in ciliogenesis [89]. Similarly, we found that several EJC core components (i.e. MAGOH, EIF4A3 and RBM8A) are partially localized to cell junction regions and that depletion of MAGOH disrupts the localized translation of polarity markers in human MCF7 cells. Moreover, through *in vivo* RNAi studies in follicular epithelial cells of the fly ovary, we show that depletion of *Mago* and *Tsu*, the fly orthologues of *Magoh* and *Y14*, disrupts localization of junctional mRNAs and proteins and impacts morphology of this tissue. By contrast, depletion of *Btz*, which encodes the orthologue of the MLN51/CASC3 protein, which is viewed as a more variable component of the EJC, resulted in a normal ovarian phenotype [85]. Previous studies have also shown that localized mRNAs that are natural NMD targets (i.e. that contain introns in their 3’UTRs), can be programmed to undergo targeted NMD following a pioneer round of translation in synaptic regions of cortical neurons or in pathfinding axonal growth cones [90, 91], a mechanism of translation-mediated mRNA decay that can fine-tune protein expression in specific subcellular compartments. While the junctional transcripts that we studied herein appear not to be natural NMD targets, suggests that NMD pathways would likely not be involved in the local turnover of the mRNA; however, it is possible that other junction targeted mRNAs could undergo translation-coupled decay. Altogether, we speculate that several of the identified cell junction RBPs may associate within regulatory modules to fine tune RNA activity at the cell periphery in order to influence epithelial cell polarity and cell-cell adhesion behavior, for example during epithelial to mesenchymal translation states or in of epithelial cancers. In this context, our findings that ZO-1 and MAGOH expression is correlated in epithelial tissues of HGS-EOC specimens, and that expression of these factors appears to predict favorable disease prognosis is quite intriguing. These data are consistent with our dissection of the positive regulatory impact of MAGOH on localized translation of *Zo-1* mRNA and in the maintenance of polarity features in epithelial cell lines. Further dissection of how impairments in localized RNA regulatory events contribute to cancer-associated loss of polarity phenotypes will be a fascinating area of future investigation.

## ACKNOWLEDGEMENTS

We thank the Bloomington *Drosophila* Stock Center for various fly stocks, especially the DRSC/TRiP project. We thank members of the Lécuyer laboratory for discussion and review of the manuscript prior to submission. We thank Xiaofeng Wang and Dominic Filion for advice on microscopy and image analysis. A.C. was funded by the Vanier Canada Graduate Scholarship program from Canadian Institute of Health Research (CIHR), as well as major scholarships from McGill University and IRCM Foundation. J.Be. is funded by the Fonds de Recherche du Québec Santé (FRQS) postdoctoral research scholarship. E.L. is an FRQS Senior research scholar. This work was supported by grants to E.L. from CIHR and the Canadian Cancer Society.

## AUTHOR CONTRIBUTIONS

A.C. and E.L. were responsible for conceptualization, methodology and investigation. A.C., J.Be. and L.C. performed experiments and/or data analyses. J.Bo. performed bioinformatics analyses.

A.C. and E.L. wrote and edited the manuscript. E.L. and A.M. supervised and/or obtained funding for the project or infrastructures employed.

## DECLARATION OF INTEREST

The authors declare no competing interests.

## MATERIAL AND METHODS

### Drosophila strains

Oregon R (OreR) was used as wildtype (WT). *TJ-Gal4* was a kindly provided by Dorothea Godt (University of Toronto). *MAGO-RNAi-TRiP* (stock 28931) and *TSU-RNAi-TRiP* (stock 28955) fly lines were obtained from the Bloomington Drosophila Stock Center (BDSC).

### Probe synthesis for FISH in *Drosophila*

Digoxigenin-labeled antisense RNA probes were synthesized as previously described [92]. Briefly, T7/Sp6 promoter-flanked cDNA templates were amplified by PCR from the *Drosophila* Gene Collection (pyd= LD02541; cno= LD24616, scrib = LD43989), following published procedures. Using 500ng of gel purified (Qiagen, 28704) and ethanol precipitated DNA templates, RNA probes were synthesized by *in vitro* transcription in the presence of Dig-UTP nucleotide labeling mix with T7 or SP6 RNA polymerases at 37°C for 4-6h. Transcription reactions were subsequently precipitated, dosed and stored at −80°C.

### FISH and IF in *Drosophila* Embryos and Egg Chambers

Embryos were harvested from population cages, and processed for IF and FISH as described previously [92]. Briefly, embryos were harvested, dechorionated with 3% bleach, fixed with 4% paraformaldehyde (PFA) in a 25% PBS and 75% heptane mixture for 20 min, transferred to a 50:50 methanol:heptane mixture to crack vitelline membranes, then stored in methanol at-20°C. Ovaries were dissected from well-fed flies using forceps in ice-cold PBS on deep-well depression slides, transferred to 1.5 mL tubes within 15 min and fixed in PBS + 4% PFA + 1% DMSO for 1h at room temperature on a nutating mixer [93]. Fixed ovaries were washed with PBS and dehydrated via successive incubations with PBS containing 25%, 50% and 75% ethanol for 5min each and then stored in 100% ethanol at-20°C.

FISH was performed as detailed previously [92, 93]. Stored embryos or dissected ovaries were rehydrated into PBS + 0.1% Tween-20 (PBTw), subjected to permeabilization steps with Proteinase K, thoroughly washed in PBTw, transferred into RNA hybridization buffer (RHB) containing 50% Formamide, 5 × SSC, 0.1% Tween-20, 100 μg/mL Heparin and 100 μg/mL salmon sperm DNA. After 2h of pre-hybridization at 56°C, RHB containing 200-400ng of DIG-labelled antisense RNA probe was added to the sample and hybridized overnight at 56°C. The next day, samples were successively washed with RHB, RHB:PBT (3:1, 1:1, 1:3) and PBT at 56°C, transferred to room temperature for 3 additional PBTw washes, then incubated with PBTw + 1% non-fat dry milk (PBTwM) for 15 min for subsequent staining with mouse Anti-Dig antibody (Dilution 1:400) for 2h in PBTwM on a nutator. After 5 × 6 min washes with PBTM, the samples were then incubated with streptavidin-HRP reagent (Dilution 1:100) in PBTM for 2h. Finally, following several PBTM washes, the samples were incubated with Tyramide-Cy3 (Dilution 1/50) from the TSA Kits (Perkin Almer) for 2h, prior to washing them in PBT and their storage in mounting solution. For IF, stored egg chambers were rehydrated into PBTw, saturated with PBTw + 2 % BSA (BBTwB) for 1h and incubated with appropriate primary antibodies overnight at 4°C on a nutator. After 5 washes with PBTwB, samples were incubated with appropriate species-specific secondary antibodies for 2h at room temperature, followed by an additional series of PBTwB washes. The DAPI staining was performed during the last PBTwB washes, after which the samples were mounted in a DABCO solution (2.5% DABCO, 70% Glycerol, 1X PBS).

### Cell Culture

MCF7 and HEPG2 cell lines were kindly provided by Brenton Graveley (UConn Health Center). MCF7 cells were cultured in growth medium containing MEM with 2mM L-glutamine (Cellgro, 10-010-CM), 1% non-essential amino acids (Cellgro, 25-025-CL), Sodium Bicarbonate 1.5g/L (Cellgro, 25-035-CL), 10% FBS (Sigma, F1051-500) and 1% penicillin/streptavidin (Wisent, 450- 201-EL). HEPG2 cells were cultured in Dulbecco’s modified Eagle’s medium (Hyclone, SH30022.01) containing additional 10% FBS and 1% penicillin/streptavidin (Multicell, 450-201EL). HMLE cells were cultured in medium containing 50% MEGM (Sigma, 815-500), 25% DMEM (Multicell, 319-005-CL) and 25% F12 (Multicell, 318-010-CL), supplemented with 10 ng/ml of hEGF (Wisent, 511-100UM), 10 ug/ml of insulin (Sigma, 10516) and 0.5 ug/ml of hydrocortisone (Sigma, H0888-1G).

### Antibodies

Our collection of rabbit polyclonal antibodies targeting human RBPs was described previously [40, 42] and contains specimens from Bethyl, Genetex and MBL. The antibodies were generally used at a final concentration of 2μg/mL in PBTB (PBS + 2% BSA + 0.3% Triton X100) during incubation. Cell junction marker antibodies included mouse anti-ZO1 (339100, Invitrogen, 1:300 dilution); rat anti-Ecad (ab11512, Abcam, 1:300 dilution); rabbit anti-SCRIB (Gtx107682, Genetex, 1:200 dilution); rabbit anti-ZO1 (Gtx108613, Genetex, 1:200 dilution); rabbit anti-AFDN (Gtx11337, Genetex, 1:200 dilution); mouse anti-DLG1 (4F3, DSHB, 1:200 dilution).

Production of a custom rabbit anti-PYD was achieved as follows. The *pyd* cDNA (LD02541) was obtained from the *Drosophila* Gene Collection and used to amplify a region encoding amino acids 1-294 of PYD. This sequence was inserted into the pET-23A (Novagen) bacterial expression vector, using EcoRI and HindIII restriction sites, in frame with a HIS-tag cassette. Following IPTG-induced expression in *e. coli* BL21 cells, bacterial cell lysates were prepared and HIS-tagged PYD(1-294) was purified on nickel-chelated agarose beads (Thermo Scientific, 78320), according to the manufacturers recommendations and dosed by Lowry assays (Bio-Rad). Protein induction and expression were verified by Coomassie gel using Brilliant Blue dye. For antibody production, purified HIS-tagged PYD was subjected to dialysis in 1L PBS overnight at 4 °C and sent to Cacolico Biologicals for rabbit immunization, for a total of 3 injections. The immune serum was collected and validated by Western blotting and IF (used at 1:100).

### Immunofluorescence and Image Analysis

After 72h of growth in a controlled incubator with 5% CO_2_ at 37 °C, cells were fixed in PBS containing 3.7% formaldehyde (Sigma, 252549) for 20 min. All steps were conducted in room temperature, unless otherwise stated. Cells were permeabilized for 20 min in PBS with 0.75% Triton X-100, followed by blocking in PBTB (PBS, 2% BSA, 0.3% Triton X-100) for 1 h. Cells were incubated with primary antibodies against cell junction markers and RBPs overnight at 4°C in PBTB according to the concentration and dilution described in previous section. Cells were washed 3 times in PBT (PBS, 0.3% Triton X-100) for 20 min each. Cells were incubated with secondary antibodies (Cy3-conjugated anti-rabbit, Jackson ImmunoResearch 311010CC; anti-rabbit Alexafluor 488 DAR, Invitrogen R37118; anti-mouse Alexafluor 647, Invitrogen A31571; anti-mouse Alexafluor 594 DAM, Invitrogen R37115; Cy5-conjugated anti-rat, Invitrogen A21434; anti-rat Alexafluor 555, Invitrogen A21434), diluted according to manufacturer’s recommended concentration in PBTB for 1 h at room temperature. Cells were labeled with DAPI (D3571, Invitrogen) for 10 min in PBTB. Finally, cells were washed 3 times in PBT for 10 min each and 3 times in PBS for 10 min each. Cells were either stored in PBS for 96-well plates or were mounted with ProLong Diamond Antifade Mountant solution (P36962, Invitrogen) for slides, which were subsequently kept at 4°C in the dark until imaging. Images from 96-well plate screens were captured by an ImageXpress Micro (Molecular Devices) high content screening microscope, while all other images were captured on a LSM700 confocal microscope (Zeiss).

### siRNA Transfections

To transfect MCF7 cells, two Silencer Select siRNA (Ambion) per target were pooled (10nM each) in Opti-MEM I medium. Lipofactamine (RNAiMAX) was added to each well according to manufacturer’s instruction. siRNA pairs with inefficient knockdown efficiencies, as revealed by subsequent RT-qPCR analyses, were omitted from further downstream assays. Cells were seeded in complete growth medium without antibiotics such that the final siRNA concentration is 20nM. Cells were then incubated at 37°C and 5% CO_2_ in environmentally controlled chamber until desired confluency, prior to downstream assays.

### Puro-PLA in MCF7 Cells

This assay was generally carried out as previously described [35], using standard 8-well chambered slides in which 100K cells were seeded per well. These assays were either performed with untreated MCF7 cells, or cells transfected with siRNAs, as detailed above. Three days post-seeding, 2μM of Puromycin (Puro) was added to the wells for 5min in full culture medium at 37 °C in a humidified atmosphere with 5% CO_2_. In parallel, as a negative control, a ‘No Puro’ samples were processed in all experiments. For the Anisomycin (Torcris) control specimens, prior to Puro treatment, cells were first pretreated with 62 μM of Anisomycin for 30min in the full culture medium at 37 °C. To arrest treatment, two quick washes were performed in pre-warmed PBS-MC (1× PBS, pH 7.4, 1 mM MgCl_2_ from Biobasic MB03228; 0.1 mM CaCl_2_ from Biobasic CD0050). Cells were fixed for 20 min in PFA-sucrose (4% formaldehyde from Sigma 252549, 4% sucrose from Biobasic SB0498, in PBS-MC) at room temperature and washed with PBS. Cells were permeabilized in PBS with 1% Triton X-100 for 15min. To reduce non-specific staining, cells were incubated in Duolink blocking solution for 1h at 37°C in humidified chamber. Afterwards, rabbit anti-SCRIB (Genetex, diluted 1:200), rabbit anti-ZO1 (Genetex, diluted 1:200) or rabbit anti-AFDN (Genetex, diluted 1:200) and mouse anti-Puro (EQ0001, Kerafast, diluted 1:2500) were used as antibody pairs for 1.5h in Duolink antibody diluent (Sigma) at room temperature. Two 5min washes with wash buffer (10 mM Tris from Biobasic TB0103, 0.15 M NaCl from Bioshop SOD001.5, 0.05% Tween-20 from Biobasic TB0560) were performed. Using Duolink antibody diluent, anti-rabbit PLAplus and anti-mouse PLAminus probes (Sigma) were diluted 1:5, and incubated at 37°C for 1h in humidified chamber. Two 5min washes with wash buffer were performed. Following the company’s recommendation, the Duolink Detection reagents Orange kit (Sigma) was used for the ligation and amplification steps, although the amplification incubation at 37°C was extended to 1h. Upon the completion of Puro-PLA assays, cells were stored overnight at 4°C in PBS. Cells were blocked in PBTB (PBS, 0.75% Triton, 1% BSA) for 1h at room temperature and were subsequently stained with anti-rabbit Alexa Fluor 488 secondary antibody (Invitrogen) for 1h in PBTB. Three 10min washes in PBS were then performed, then samples were mounted with Duolink mounting solution containing DAPI and coverslips were sealed with nail polish. Samples were imaged with the Zeiss LSM 700 confocal system with a 63x oil objective and a pinhole setting of 1AU.

### RNA Immunoprecipitation (RIP)

To prepare cell lysates for RIP, cells were harvested from cultures at 70-95% confluency, washed twice with ice cold PBS, scraped off the plates and spun at 2000g at 4°C for 5 mins. Cell pellets were resuspended in NP-40 lysis buffer (150mM NaCl, 1% NP-40, 50mM Tris pH 8.0) with protease inhibitors (Aproptinin, Leupeptin, Pepstatin A, PMSF, Biobasic) and RNAseOUT (Invitrogen) at 4°C. Cells were then rocked on ice at 4°C for 10mins, centrifuged at 720g for 5min at 4°C to eliminate nuclei. The supernatant containing the cytoplasmic and membrane compartments was recovered and dosed via Lowry assays (Bio-Rad 5000113, 5000114, 5000115). Extracts were then flash frozen in liquid nitrogen and stored at-80°C.

In preparation for immunoprecipitation (IP), primary antibodies (10 µg) were cross-linked to magnetic protein A/G beads (Thermoscientific) in freshly prepared solution containing 0.2M triethanolamine pH 8.2 diluted in PBS and 20mM DMP (Thermoscientific, PI-21667) by incubating for 30 min at room temperature. Primary antibodies used include anti-PCBP3 (Mbl, RN054PW); anti-MAGOH (Mbl, RN089PW); anti-SNRNP200 (Mbl, RN095PW) and anti-PRPF4 (Mbl, RN093PW), while rabbit IgG (Sigma) was used in parallel as a specificity control. To begin IP assays, cellular extracts were first pre-cleared by incubation with freshly prepared protein A/G beads in binding buffer (50mM Tris pH 7.4, 150mM NaCl, 1mM MgCl_2,_ 0.05% NP-40) for 1h at 4°C on a nutator. Pre-cleared cell lysates (1mg per IP sample) were then recovered and incubated for overnight at 4°C on nutator with 30 μL of crossllinked antibody-bead conjugate (∼10 µg of antibody/sample) in a total volume of 1mL of binding buffer containing 250U RnaseOUT and 0.5 mM fresh DTT, while setting aside 1% of the sample as input. Antibody-conjugated AG beads were then captured on magnetic columns, washed 6 times with 1 mL binding buffer and resuspended in 500 µL of Proteinase K buffer (10% SDS, 50 mM EDTA, 10 mM Tris-Cl pH 7.4, 25mg/mL Proteinase K, Sigma P2308) and incubated at 50°C for 45 min with agitation, followed by cooling on ice. Input samples were treated with Proteinase K buffer in parallel. To extract the RNA, 1mL TRIzol was added and the samples were froze in liquid nitrogen and transferred to storage at-80°C. Upon retrieval, chloroform (VWR Analytical BDH1109-4LG) was added to TRIzol-treated sample, vortexed and centrifuged at 14000g at 4°C. To isolate the RNA in the aqueous phase, the top layer was collected and glycogen was added to assist with precipitation (Thermo Scientific R0551). Following, 0.5 volume of 100% isopropanol (Fisher Bioreagents BP2618-4) was added, incubated and centrifuged at 14 000 g at 4°C, followed by centrifugation with 80% EtOH at 4°C. RNA pellet was dried and resuspended in small volume of water, which was further saved at-80°C until required for cDNA synthesis.

### cDNA Synthesis and RT-qPCR

Using TRIzol reagent, total RNA was isolated from cells as described by manufacturer’s protocol (Ambion, 15596018). For each siRNA-treated vs control sample, equal amount of RNAs (1ug) were reverse-transcribed to DNA using DNAseI (NEB, M03035) for cleanup, EDTA for inactivation of reaction, and finally, random hexamers (Promega C118A) and the M-MLV reverse transcriptase kit (Invitrogen, 28025-013) for cDNA synthesis of each sample. For each RNA immunoprecipitation example, an equal volume of RNAs were used as cDNA template for reverse transcription instead. qRT-PCR was conducted with the PowerUp SYBR Green Master Mix (Applied Biosystems, 100029283) using gene-specific primer pairs **(Table.S2, S3)** on an Applied Biosystems ViiA7 Real-time PCR instrument. The relative expression for each sample was computed using the Ct method through comparison to internal housekeeping genes such as *Rps16*, *Ppia*, or *Gapdh* **(Table.S2, S3)**.

### Tissue Microarray

Ethics and materials transfer approval was obtained from the Centre Hospitalier de l’Université de Montréal (CHUM) institutional ethics committee. Informed patient consent was obtained. HGS carcinoma tissues were extracted during primary cytoreductive surgery of patients, which was subsequently formalin-fixed and paraffin-embedded (FFPE). In this study, we used a discovery cohort, consisting of HCS-EOC tissues from 101 patients, which were recruited at CHUM between 1993 and 2012. The tumour staging and histotype were graded by a pathologist using standards set by the International Federation of Gynecology and Obstetrics (FIGO). IF on TMA slides was performed on 4uM TMA tissues through pipetting and use of the Benchmark XT autostainer (Ventana Medical Systems, Roche). The epithelium was identified by a co-staining of epithelial cytokeratins using a cocktail of mouse antibodies against KRT7 (Neomarkers, MS-1352- P), KRT18 (Santa Cruz Biotechnology, sc-6259), and KRT19 (Thermo Scientific, MS-198-P) at 1:200. MAGOH was detected using a rabbit antibody (Mbl, RN089PW) at 1:200 and ZO-1 was detected using a mouse antibody (Invitrogen, 339100) at 1:200. Slides were then incubated with corresponding secondary fluorescent antibodies (Cyanine 7 for cytokeratin, Cyanine 5 for MAGOH and AF488 for ZO-1 detection) for 45 min. DAPI was used to stain the cell nuclei. Samples were mounted on slides using the Fluoromount medium (Millipore-Sigma, F4680). Computerized image analysis was performed as previously described using the Visiopharm software (Hoersholm) and OlyVIA image viewer software (Olympus) [44]. Briefly, DAPI was used to mark the nuclei, cytokeratins were used to differentiate region of interest, such as epithelium from stroma in each specimen. Protein expression was measured in each image pixel of each region of interest to obtain MFI. A visual review was performed to exclude experimentally damaged tissue and inflammatory sections, which may boost artificial signal of markers.

Statistical analyses were conducted using IBM Statistics SPSS 23. Correlation analyses between markers were performed using non-parametric Spearman correlation. The non-parametric Mann-Whitney test was used to compare the mean between two groups. Multiple group comparison was performed using the Anova test followed by the post-hoc Tuckey test. For mean comparisons, FIGO stages were dichotomized into groups of early stages (1 and 2) versus late stages (3 and 4). MAGOH expression was categorized into four equal groups of patients by MFI expression quartiles using cut-offs at the 25th, 50th and 75th percentiles. Kaplan–Meier curves were performed to dissect marker association with patient prognosis. For survival analysis, MAGOH expression was dichotomized into groups of negative/low and medium/high expression by the 25th percentile of MFI. A p-value < 0.05 was considered statistically significant (* = p < 0.05; ** = p < 0.01; *** = p < 0.001).

### Microscopy

Samples were imaged on a Zeiss LSM700 laser scanning confocal microscope with a pinhole setting of 1AU, using a 40x or 63x oil objective, unless otherwise stated. For high-throughput experiments in 96-well plates (Corning 3904, Corning 3882) images were captured by the ImageXpress Micro high content screening system made by Molecular Devices lnc., using a 40x objective. In order to capture the best intensity, an auto exposure function was used. The exposure time for the RBP channel, marker channel and DAPI ranged from 250-5000, 250-3000 and 100-1000 ms, respectively. The built-in laser-based auto-focusing function was utilized to achieve the best image focus in a rapid and automated fashion. High content images were further analyzed using the MetaXpress v3.1 software. TMA slides were scanned with the VS-110 microscope using 0.75 numerical aperture with a 20X objective. Figures were constructed using Adobe Photoshop and Illustrator.

### Quantification and statistical analysis

Specific information on statistical tests is explained in the respective figure legends. Statistical analyses were performed using GraphPad Prism 8 software. P values and error bars are indicated in the relevant graphs.

### eCLIP database survey

To elucidate whether there are direct binding peaks between candidate RBPs and cell junction mRNAs of interest, we conducted a survey of different CLIP-seq datasets on the CLIPdb database [43]. Notably, data were available for eight RBP hits, in HEPG2 or HEK293 cells.

## Notes

### Competing Interest Statement

The authors have declared no competing interest.

