## Supplement Figures and Tables for "Localized translation of cell junction mRNAs is required for epithelial cell polarity"

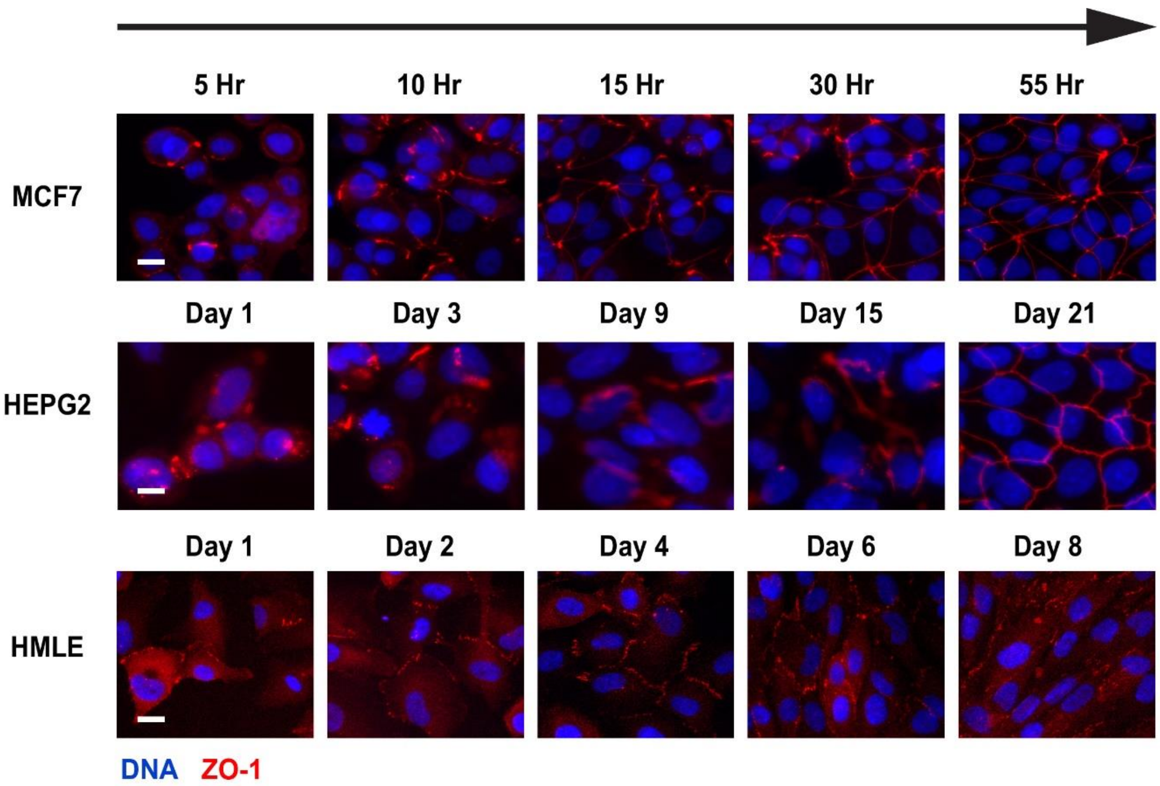


**Figure S1. Epithelial cell lines display various rates of cell junction marker localization.**

Temporal IF over 5 timepoints post-seeding of MCF7, HEPG2 or HMLE cells. Localization of ZO-1 protein (red) is used as a common marker of cell-cell junction regions. Blue, DNA. Scale bar represents 15µm.

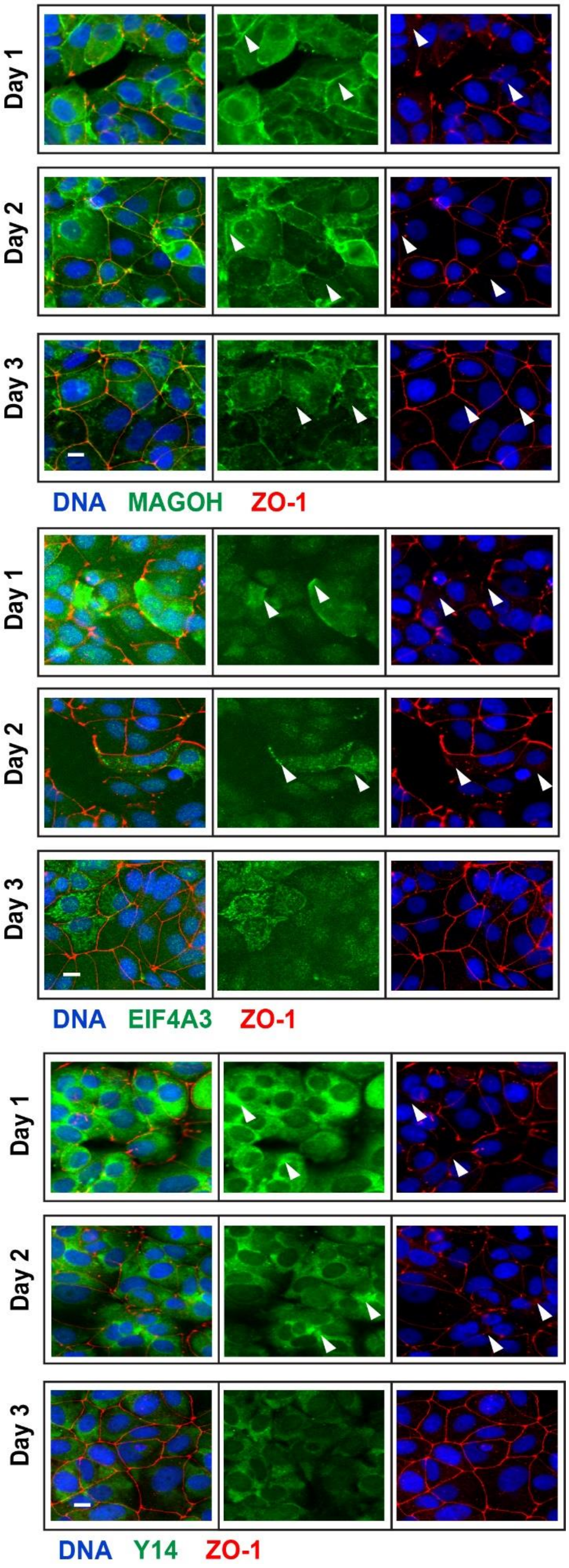

Day 1
Day 2
Day 3
DNA MAGOH ZO-1
Day 1
Day 2
Day 3
DNA EIF4A3 ZO-1
Day 1
Day 2
Day 3
DNA Y14 ZO-1

**Figure S2. Core components of EJC localize to cell-cell junction regions.**

Temporal, IF-based localization study of the core components of the EJC, including MAGOH, EIF4A3 and Y14, in MCF7 cells. ZO-1 is used as a marker of cell-cell junctions. Arrowheads indicate sites of enriched localization of EJC component at cell-cell borders, which often appeared to be targeted to the emerging cell junction region prior to ZO-1, especially in day 1 and 2. By day 3, the expression of EJC components begin to diminish. Red, ZO-1 protein. Green, protein. Blue, DNA. Scale bar represents 15µm.

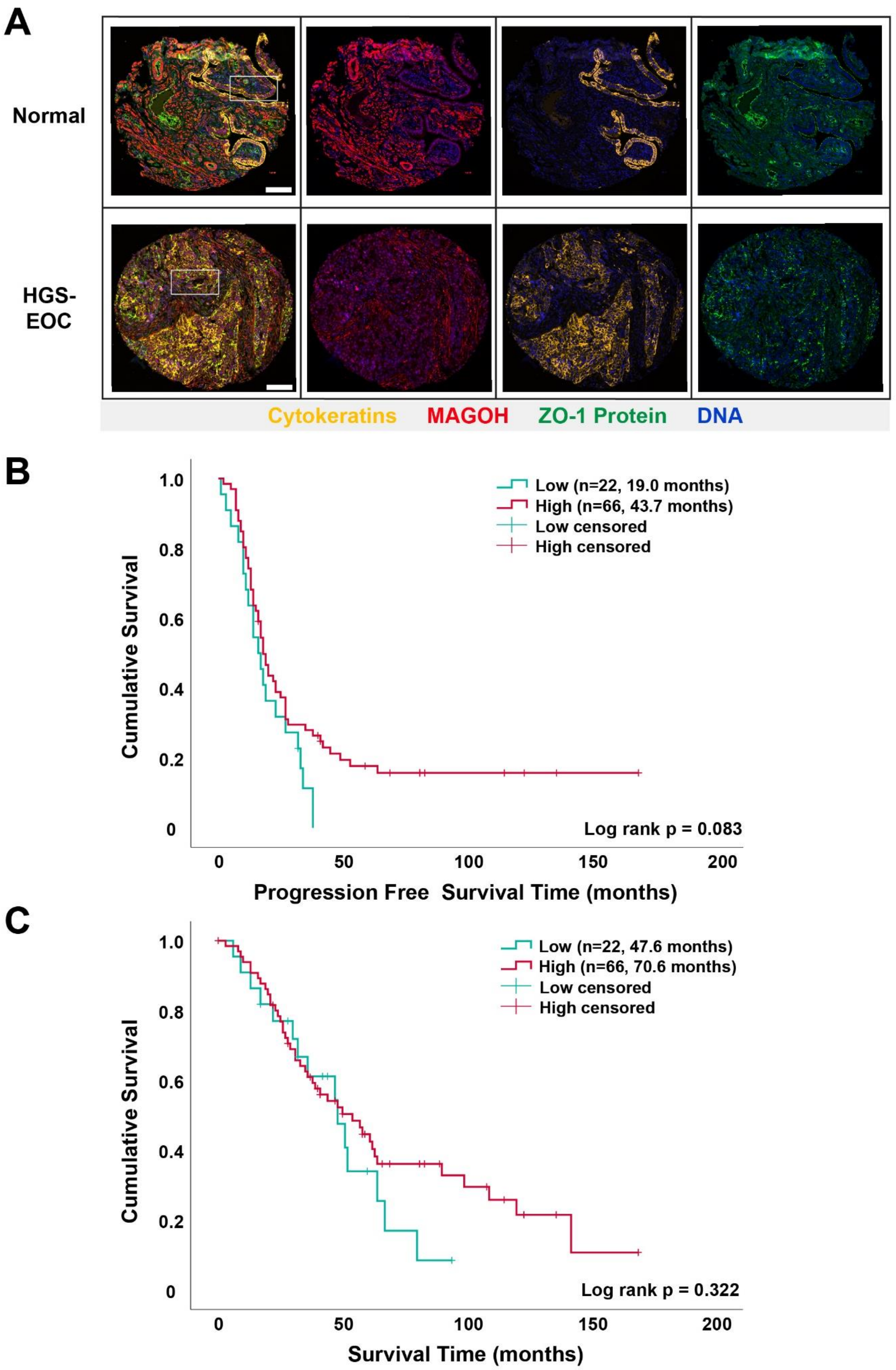

A
Normal
HGS-
EOC
Cytokeratins
MAGOH
ZO-1 Protein
DNA
B
Low (n=22, 19.0 months)
High (n=66, 43.7 months)
Low censored
High censored
Cumulative Survival
1.0
0.8
0.6
0.4
0.2
0
Log rank p = 0.083
0
50
100
150
200
Progression Free Survival Time (months)
C
Low (n=22, 47.6 months)
High (n=66, 70.6 months)
Low censored
High censored
Cumulative Survival
1.0
0.8
0.6
0.4
0.2
0
Log rank p = 0.322
0
50
100
150
200
Survival Time (months)

**Figure S3. Visualization of ZO-1 and MAGOH protein expression in tissue microarrays of a discovery cohort of ovarian cancer specimens.**

**(A)** Examples of IF results obtained for ZO-1 and MAGOH proteins on TMA panels consisting of surgically extracted human ovarian tissues, including normal fallopian tubes (n=15) and HGS-EOC tumor specimens (n=101) **.** Normal tissues reveal that MAGOH expression is expressed in both epithelium and surrounding stroma, with organized Cytokeratins and ZO-1 staining at the highly polarized epithelium. However, a general disorganization of these three markers is observed in HCS. White squares denote the enlarged view of the selected region shown in **Figure 7D**. DAPI (blue), MAGOH (red), Cytokeratins (yellow), ZO-1 (green) and DAPI staining are shown. Scale bars represent 100µM. **(B)** Kaplan-Meier curves of MAGOH expression and its association with progression-free survival. MFI was used as measure of expression for markers, which was dichotomized by the median for MAGOH by the 25$^{th}$ percentile cutoff, into groups belonging to low (n = 22) or high expression (n = 66). P-value of 0.083 was observed by Mantel-Cox analysis. **(C)** Kaplan-Meier curves of MAGOH expression and its association with survival time. MFI was quantified for markers, which was separated into groups as above. P-value of 0.322 was observed by Mantel-Cox analysis.

**Table. S1. General information on candidates identified from RBP-focused screen which exhibited cell junction-like distribution patterns.** Table detailing description of proteins identified, signal strength and localization attributes in MCF7.

| Mammalian RBP | Description | Junctional Signal | Tri-cellular Pattern | Nuclear Localization | Cytoplasmic Localization |
|---|---|---|---|---|---|
| WDR3 | WD repeat domain | Strong | Yes | Yes | No |
| DDX20 | DEAD box polypeptide | Weak | Yes | Yes | Yes |
| GNL3 | Guanine nucleotide binding protein-like | Weak | Yes | Yes | Yes |
| CORO1A | Coronin, actin binding protein | Weak | Yes | Yes | Yes |
| KIF1C | Kinesin family member | Weak | Yes | Yes | Yes |
| TRIP6 | Thyroid hormone receptor interactor | Strong | Yes | Yes | Yes |
| SMARCA3 | Helicase-like transcription factor | Weak | No | Yes | Yes |
| NELF-E | Negative elongation factor complex | Strong | No | Yes | Yes |
| NPM1 | Nucleophosmin | Strong | Yes | Yes | No |
| MAGOH | Component of EJC | Strong | No | Yes | Yes |
| PCBP3 | Poly(RC) binding protein | Strong | No | Yes | Yes |
| BTF | BCL2-associated transcription factor | Weak | Yes | Yes | No |
| DGCR8 | Microprocessor complex subunit | Strong | Yes | Yes | Yes |
| ATXN1 | Spinocerebellar ataxia type protein | Weak | No | Yes | Yes |
| SNRNP200 | Small nuclear ribonucleoprotein | Weak | No | Yes | Yes |
| PRPF4 | Pre-mRNA processing factor | Weak | No | Yes | Yes |
| PRKCBP1 | Zinc finger, MYND-type | Strong | Yes | Yes | Yes |
| DDX1 | DEAD box helicase | Weak | Yes | Yes | Yes |
| KRR1 | Small subunit processome component | Weak | No | Yes | Yes |
| SSRP1 | Structure specific recognition protein | Strong | No | Yes | Yes |
| DNAJC17 | DnaJ homolog | Weak | No | Yes | Yes |
| CSTF2 | Cleavage stimulation factor subunit | Weak | No | Yes | Yes |
| RBFOX2 | Fox-1 homolog | Weak | No | Yes | Yes |
| MPHOSPH6 | M-phase phosphoprotein | Strong | No | Yes | Yes |
| THUMPD2 | THUMP domain containing protein | Weak | No | Yes | Yes |

**Table. S2. Primers used for RNA immunoprecipitation.** A list showing the multiple primer sets utilized for testing association of RBP candidates with selected cell junction mRNAs when compared to housekeeping controls.

| **Primer ID** | **Sequence** |
|---|---|
| Zo1_1_Fw | ggaggtagaacgaggcatc |
| Zo1_1_Rv | gagcggacaaatcctctctg |
| Zo1_2_Fw | ttgcaatatctggtggacg |
| Zo1_2_Rv | tgaaactccgttaaccattgc |
| Zo1_3_Fw | aaagagatgaacgggctacg |
| Zo1_3_Rv | ggggaggcctatcgtgtg |
| Scrib_1_Fw | agctgatcctcacggagaac |
| Scrib_1_Rv | ctcaaggagaggacgctgag |
| Scrib_2_Fw | agcccctgaagctggactac |
| Scrib_2_Rv | ctaggggagcagggagattc |
| Scrib_3_Fw | ctcactgacctgctgctgtc |
| Scrib_3_Rv | aggttctccgtgaggatcag |
| Dlg1_1_Fw | caacccacacattggagatg |
| Dlg1_1_Rv | cctgcttctttcaacgcttc |
| Dlg1_2_Fw | cgttaatggcacagatgcag |
| Dlg1_2_Rv | catctccaatgtgtgggttg |
| Dlg1_3_Fw | aacgagcccgattaaaaac |
| Dlg1_3_Rv | ttcactatcgctggcattag |
| Afdn_1_Fw | aatgggccttagcattgttg |
| Afdn_1_Rv | accagacttcgtccatccac |
| Afdn_2_Fw | gaatgctgtcctctcccaag |
| Afdn_2_Rv | ttcccgatcatctttgttcc |
| Afdn_3_Fw | caaagctttaccgccttcag |
| Afdn_3_Rv | tcacagtgaccactccatcc |
| Rps16_1_Fw | tgcagtctgtgcaggtcttc |
| Rps16_1_Rv | cgttcaccttgatgagacc |
| Rps16_2_Fw | ggcaatggtctcatcaagg |
| Rps16_2_Rv | accaccctttacacggacac |
| Rps16_3_Fw | tggtgtagacatccgtgtcc |

| | |
|---|---|
| Rps16_3_Rv | ttcttggaagcctcatccac |
| Ppia_1_Fw | cctaaagcatacgggtcctg |
| Ppia_1_Rv | ggcctccacaatattcatgc |
| Ppia_2_Fw | agggttcctgctttcacag |
| Ppia_2_Rv | caggacccgtatgctttagg |
| Ppia_3_Fw | ggcctggataccaagaagtg |
| Ppia_3_Rv | gtctttgggaccttgtctgc |

**Table. S3. Primers used for testing siRNA efficiency.** A list showing the primer sequences used for testing knockdown of RBP candidates.

| Primer ID | Sequence |
|---|---|
| WDR3_Fw | ggggttggttctgttgtcag |
| WDR3_Rv | ttgattttcttggggtctgg |
| DDX20_Fw | tacattggggctgacagtga |
| DDX20_Rv | cccaatccacacattcttcc |
| GNL3_Fw | gccagagatcctcttggttg |
| GNL3_Rv | tgaggctctgaacaccactg |
| CORO1A_Fw | ccttctttgaccctgacacc |
| CORO1A_Rv | tggcgatctcacacttgttc |
| KIF1C_Fw | tcacccaccacacataatgg |
| KIF1C_Rv | tctcctcccatgtctcgttc |
| HLTF_Fw | accagtgaaaaggcagatgg |
| HLTF_Rv | gcgggagctagacaattctg |
| NELF-E_Fw | ccgttccagaggagcatatc |
| NELF-E_Rv | ttcatagccccagtcaaagc |
| NPM1_Fw | ttgttgaagcagaggcaatg |
| NPM1_Rv | aatatgcactggccctgaac |
| BTF_Fw | agctgcaatgaccctaaacg |
| BTF_Rv | tgctaaacgggtatgcttcc |
| ATXN-1_Fw | gtgaagcctccccttctacc |
| ATXN-1_Rv | gtggtctgaatgaccgtgtg |
| SNRNP200_Fw | agaagctggagctgcagaag |
| SNRNP200_Rv | tgtgagatgaaggcttgcag |
| PRPF4_Fw | agtcagattggggatgatcg |
| PRPF4_Rv | tgcaatcaggaacagaccag |
| DDX1_Fw | cagactctgggaaaggcttg |
| DDX1_Rv | cagcatactccccctttcagg |

| | |
|---|---|
| KRR1_Fw | tttggaggagagcagtttcg |
| KRR1_Rv | caaacagtcatgctgccttc |
| DNAJC17_Fw | ccacccagacaaaaatccag |
| DNAJC17_Rv | cttgcttcttggctttcctg |
| RBFOX2_Fw | aagcccagtagttggagctg |
| RBFOX2_Rv | tcttcctgataggggcactg |
| THUMPD2_Fw | ccaggaagttggttgaatgc |
| THUMPD2_Rv | cagctggcagtccctattc |
| TRIP6_Fw | atgagctggataggctgacg |
| TRIP6_Rv | atccccaaccacatcttctc |
| MAGOH_Fw | aaagcgtgatggaggaact |
| MAGOH_Rv | ccttctggatccttggattg |
| PCBP3_Fw | atttgaccaagctccaccag |
| PCBP3_Rv | gactggcccatcagattttg |
| DGCR8_Fw | catcctgcacgagtacatgc |
| DGCR8_Rv | tcagggatgaggatttccag |
| SSRP1_Fw | gccaaactcgctaccacttc |
| SSRP1_Rv | atcttgcggtttaccagtgc |
| MPHOSPH6_Fw | gccagaaagagagaccatgc |
| MPHOSPH6_Rv | cctggggctttaagaacatc |
| GAPDH_Fw | atgttcgtcatgggtgtg |
| GAPDH_Rv | ggtgctaagcagttggtgg |
